# Circulating extracellular vesicles in plasma carry accessible molecular signatures of aging in mice

**DOI:** 10.64898/2026.07.10.737625

**Authors:** Kristine A. Tsantilas, Michael Riffle, Gennifer E. Merrihew, Christine C. Wu, Gregory R. Keele, Aaron Maurais, Richard S. Johnson, Alison Luciano, Laura Robinson, Gary A. Churchill, Michael J. MacCoss

## Abstract

Cells release membrane-bound extracellular vesicles into the bloodstream laden with proteins that may reflect their physiological state. How this circulating EV proteome changes across life remains poorly understood. Identifying molecular signatures of aging in accessible biofluids could facilitate earlier intervention and monitoring of age-related disease. Many circulating aging proteome studies rely on affinity-based platforms which suffer from poor cross-species translation, ambiguous signal attribution, and inconsistent agreement between platforms. Here, we present a characterization of the aging plasma EV proteome from a cross-sectional cohort of 86 male and female C57BL/6J mice (5-31 months). We leveraged a species-agnostic EV enrichment (Mag-Net) and mass spectrometry to detect 2,575 protein groups from 15,969 peptides. Protein abundance heterogeneity increased with age and the abundance of 272 proteins were significantly correlated with chronological age including established senescence and frailty markers. Proteins increasing with age were enriched in genome maintenance pathways, while those decreasing were associated with the extracellular matrix organization and lipid metabolism. Notably, several of the strongest age-increased proteins converged on Alzheimer’s and Parkinson’s disease pathology. We observed sexual divergence in the aging EV proteome not previously characterized at this resolution. A proteomic clock built from this data accurately predicts chronological age, and peptide-level analysis reveals aging signals invisible at protein-level. These findings demonstrate that EV-enriched plasma proteomics can identify known aging markers, reveal novel sex-specific age-related changes, and generate predictive models of chronological age. This study provides a species-agnostic foundation for proteomic clocks that complement epigenetic approaches to monitor aging and evaluate healthspan.

## Introduction

Aging is a ubiquitous and heterogeneous process that impacts species across the tree of life. Along with aging comes co-morbidities that differ in timing, scope, and severity among individuals and whose prevalence often differs across species. In humans, aging and its associated co-morbidities are a threat to healthspan and lead to disability or death (Olshansky, 2018). Established hallmarks of aging are broadly conserved across species (López-Otín et al., 2023). Targeting conserved aging phenotypes has improved our understanding of age-related disease pathogenesis and generated leads to pursue novel treatments to increase healthspan and longevity such as rapamycin and metformin (Thanapairoje et al., 2023). The identification of molecular signatures of aging has emerged as a central objective of modern biogerontology, driven by the promise that tracking biological age could reveal mechanisms of age-related disease and guide intervention (Kroemer et al., 2025). Much of this effort has focused on epigenetic clocks which are computational models trained on DNA methylation patterns that predict chronological age with remarkable accuracy (Horvath & Raj, 2018). Yet the proteome, as the functional endpoint of both genome and epigenome, may more directly reflect the physiological state of an individual and extract more actionable signatures of aging.

Efforts have been made to characterize protein changes across aging tissues, including in humans (Ding et al., 2025) and mice (Keele et al., 2023, 2025; Takasugi et al., 2024). The circulating proteome has also been a recent area of focus (Lehallier et al., 2019, 2020; Oh et al., 2023; Schaum et al., 2020; Sebastiani et al., 2017; Shen et al., 2024; Siino et al., 2022), with studies that leveraged computational approaches like those used in epigenetic clocks (Lehallier et al., 2019; Oh et al., 2023). Unfortunately, many protein-level studies have relied exclusively on commercial affinity reagents without subsequent validation with a secondary method. Affinity reagents that bind human proteins often lose sensitivity in model organisms due to protein sequence changes common between species. Different aptamer platform findings often do not correlate well with each other (Hoofnagle & MacCoss, 2025; Rooney et al., 2025) and may not identify proteins unambiguously. Furthermore, measurements of clinically relevant proteins by aptamers can have poor correlation with clinical data (Baird & Hoofnagle, 2017; Faquih et al., 2021; Lopez et al., 2024). Liquid chromatography coupled to mass spectrometry (LC-MS) offers a species-agnostic and unambiguous alternative to affinity-based platforms for evaluating age-related changes in the proteome.

The plasma proteome is a particularly attractive substrate. It represents an easily accessible, minimally invasive way to acquire a snapshot of an individual’s physiology, containing proteins from throughout the organism (Geyer et al., 2017). Despite its many positive attributes, the substantial proteome dynamic range of plasma often necessitates depletion of abundant proteins or affinity reagents to enrich specific analytes (Zubarev, 2013). Such approaches are either unavailable for model organisms or lack equivalent efficacy across species. To circumvent these limitations, we used Mag-Net, an approach to enrich for circulating extracellular vesicles (EVs) (Wu et al., 2025). Much of the biology observed in the plasma proteome is derived from and trafficked throughout the body by EVs of diverse type, size, and origin (Andaloussi et al., 2013; Cocucci & Meldolesi, 2015; Meldolesi, 2018). Aging modifiers have been found in EVs and some can be isolated from one species and administered to another to impact health (Horvath et al., 2023; Lee et al., 2025; Yin et al., 2021; Yoshida et al., 2019). The Mag-Net method is species-agnostic, as isolation occurs based on the size and negative charge of EV plasma membranes. With LC-MS, this approach circumvents challenges with dynamic range, loss of translation between species, and ambiguous signal attribution, while improving the depth of proteome coverage that impacts many existing studies.

The diet, genetics, and environment of a laboratory mouse are tightly controlled, and their lifespan is thoroughly characterized (Yuan et al., 2009). Additionally, they have been a gold standard of vertebrate aging on a more compressed timescale than humans (Dutta & Sengupta, 2016). Thus, we sought to measure the aging proteome of circulating plasma EVs from C57BL/6J mice using tandem LC-MS in a cross-sectional cohort to determine whether we can identify a protein-level signature of aging and evaluate the utility of this approach as a species-agnostic platform for aging biomarker discovery.

## Materials and Methods

Detailed methods for all sections are included in the Supplementary Materials and Methods section of the Supplementary Material.

### Animal Care and Housing

C57BL/6J mice of both sexes were sourced from and housed at The Jackson Laboratory (Bar Harbor, ME, USA; Stock Number 000664) for the aging cohort. All experiments complied with the National Institutes of Health Guide for the Care and Use of Laboratory Animals (National Research Council), were approved by The Jackson Laboratory’s Animal Care and Use Committee, and were performed in accordance with the ARRIVE guidelines.

### Plasma collection

Blood was drawn using a submental collection protocol, transferred into EDTA collection tubes, and plasma isolated (14,000 rpm spin, 4°C, 10 minutes). Plasma was separated from the buffy coat, aliquoted, frozen, and stored at -80°C until processing.

### Quality control measures evaluating sample preparation, liquid chromatography-mass spectrometry system suitability, and quantitative results

To account for sample preparation variation and track system performance, we employed a quality control (QC) framework which has been described in detail previously (Tsantilas et al., 2024). This included external QC samples, internal QCs, and system suitability runs interspersed among sample runs. The composition, generation, use, and analysis of these external QC samples are in the Supplementary Methods. Plate layout used a balanced block design considering age, sex, and group size.

### Extracellular vesicle enrichment (Mag-Net), protein digestion, and clean-up

We enriched for circulating plasma EVs using the Mag-Net protocol (Wu et al., 2025) from 20 μL of plasma per individual on a Thermo KingFisher Apex including the particle enrichment, washes, clean-up, and digestion steps.

### Liquid-chromatography and mass spectrometry (LC-MS)

Liquid chromatography coupled to data independent acquisition (DIA)-MS was leveraged for peptide detection in neat plasma and Mag-Net enriched EV fraction samples. The enrichment and depletion experiment was run on a Thermo EASY-nLC 1200 coupled to a Thermo Orbitrap Exploris 480. The aging cohort analysis of the Mag-Net fraction was collected using a Thermo Vanquish Neo coupled to a Thermo Orbitrap Astral. In both cases, we collected gas-fractionated, narrow window runs of sample pools alongside the experimental samples for subsequent searches. System suitability was performed using targeted parallel reaction monitoring (PRM) runs before, during, and after experimental sample to assess the LC-MS system suitability.

### Mass spectrometry signal processing

DIA-MS data were processed using a Nextflow pipeline (https://github.com/mriffle/nf-skyline-dia-ms) with a *Mus musculus* reference proteome FASTA file (Uniprot Proteome ID: UP000000589, downloaded February 2, 2023) appended with the internal QC yeast enolase 1. All data was imported into Skyline (version 24.04.4).

### Enrichment and depletion relative to total plasma

Commercial pooled C57BL/6N mouse plasma was processed neat and with Mag-Net in parallel to evaluate the enrichment of EV proteins or the depletion of high abundance proteins in Mag-Net relative to neat plasma.

### Particle counting

We demonstrated that Mag-Net isolates membrane-bound particles using a Nanosight NS300 (Malvern Panalytical Ltd) fitted with a standard gasket (NTA4027) and using Nanoparticle Tracking Analysis (NTA) software (Malvern Panalytical Ltd, version NTA 3.4, Build 3.4.4). EVs were eluted from SAX beads in Mag-Net with 25 mM Bis Tris Propane (pH = 6.5), 1 M NaCl, 0.1% Tween 20 instead protein digestion. The particles were diluted 1:25 in water and measured at 25.4 °C ± 0.2 with five replicate particle counts.

### General peptide and protein-level analysis

Data analysis outside of Nextflow and Skyline was done using R (R version 4.4.1, http://www.r-project.org) or Python (Van Rossum & Drake, 2009). Log-transformed median-normalized peak areas were used for analyses of mass spectrometry results performed outside of Skyline unless specifically noted otherwise. All p-values have been adjusted for multiple hypothesis testing correction (p_adjusted_) using the Benjamini-Hochberg procedure (Benjamini & Hochberg, 1995) unless noted otherwise.

### Spearman correlation and Reactome pathway analysis

Spearman correlation was performed assuming a two-sided test using the age of the mice and the normalized protein abundance to calculate spearman rho (ρ) and an adjusted p-value. Using the list of proteins positively or negatively correlated proteins by Spearman Correlation (ρ > 0.3 or ρ < -0.3, p_adjusted_ < 0.05), a high-level analysis of implicated pathways was generated in R using packages in the Bioconductor framework. The associated mouse Uniprot Accession numbers were converted into a list of mouse ENTREZ gene IDs. The Reactome Pathway Enrichment was performed using the “ReactomePA” package (Yu & He, 2016) with the *Mus musculus* organism database.

### Ordinary Least Squares (OLS) Analysis

OLS analyses were implemented in Python. Raw abundances were recovered by exponentiating log2-transformed intensities, then normalized via natural-log transformation and z-score standardization. For each feature, an OLS model was fit regressing scaled abundance on age and sex using the statsmodels formula API. P-values for the age coefficient were corrected for multiple testing using the Benjamini–Hochberg false discovery rate (FDR) procedure, with features having FDR-adjusted q-values < 0.01 considered significant.

### Elastic Net Regularized Linear Regression

An Elastic Net regression model (Zou, H. & Hastie 2005) was used to predict chronological age from feature abundances. Elastic Net combines L1 (lasso) and L2 (ridge) penalties, enabling simultaneous feature selection and coefficient shrinkage (hyperparameters *α*=0.01 and *l1_ratio=0.3)*. Optimal hyperparameters were estimated using Bayesian optimization. Features were log10-transformed including sex as a binary covariate. Model performance was estimated using repeated *k*-fold cross-validation (10 folds, 5 repeats) implemented with RepeatedKFold. To prevent data leakage, z-score standardization (StandardScaler) was fit on the training partition and applied to the test partition independently within each fold. Model accuracy was assessed by mean absolute error (MAE) and *R^2^* computed via linear regression of predicted versus true values (linregress). Robust feature importances were derived by averaging Elastic Net coefficients across all folds and recording the frequency where each coefficient was non-zero. A final model was constructed using Elastic Net with all features using the same hyperparameters as during cross-validation, and these final coefficients were used to report the number of features kept or removed from the model. All analyses were implemented in Python using scikit-learn, NumPy, and SciPy.

### Declaration of generative AI and AI-assisted technologies

During the preparation of this work, Claude (Anthropic) was used for three purposes: to assist the author’s in writing portions of the analysis, to write and revise code for figure generation, and to improve the readability and conciseness of the text. In all cases the authors reviewed, edited, and tested the output, and take full responsibility for the content of the publication and for the correctness of the released code.

## Results

### Species applicability of Mag-Net and LC-MS system

Although the Mag-Net assay was developed and validated in human plasma (Wu MacCoss 2025 Nat Comm), it does not rely on affinity reagents and therefore has broad applicability across species. We confirmed this in neat mouse plasma and the Mag-Net enriched fraction. The Mag-Net fraction was enriched for proteins known to be trafficked in EVs, including CD9, CD81, CD44, PDCD6IP, and NCAM1 (Andaloussi et al., 2013; Cocucci & Meldolesi, 2015; Meldolesi, 2018), and depleted high-abundance plasma proteins, in line with our prior finding in human plasma (Wu et al., 2025). The distribution of captured particle sizes (diameter 75-250 nm) was representative of exosomes and microvesicles (Andaloussi et al., 2013; Cocucci & Meldolesi, 2015).

### Aging cohort design, proteome depth, and data quality control

We leveraged Mag-Net and DIA-MS to characterize the mouse plasma EV proteome across the natural lifespan of adult male and female C57BL/6J mice (Figure 1A). This cross-sectional aging cohort included 86 mice across 9 age groups spanning 25–131 weeks (5.9–30.3 months) with 3–6 mice per sex per group (Figure 1B). Two types of external QC samples were prepared and analyzed alongside the cohort (Figure 1C) to evaluate sample preparation consistency and data normalization: an inter-experiment QC (IE-QC) from a commercial plasma pool and an inter-batch QC (IB-QC) pooled from mice housed alongside the aging cohort (Figure 1D).

**Figure 1:**
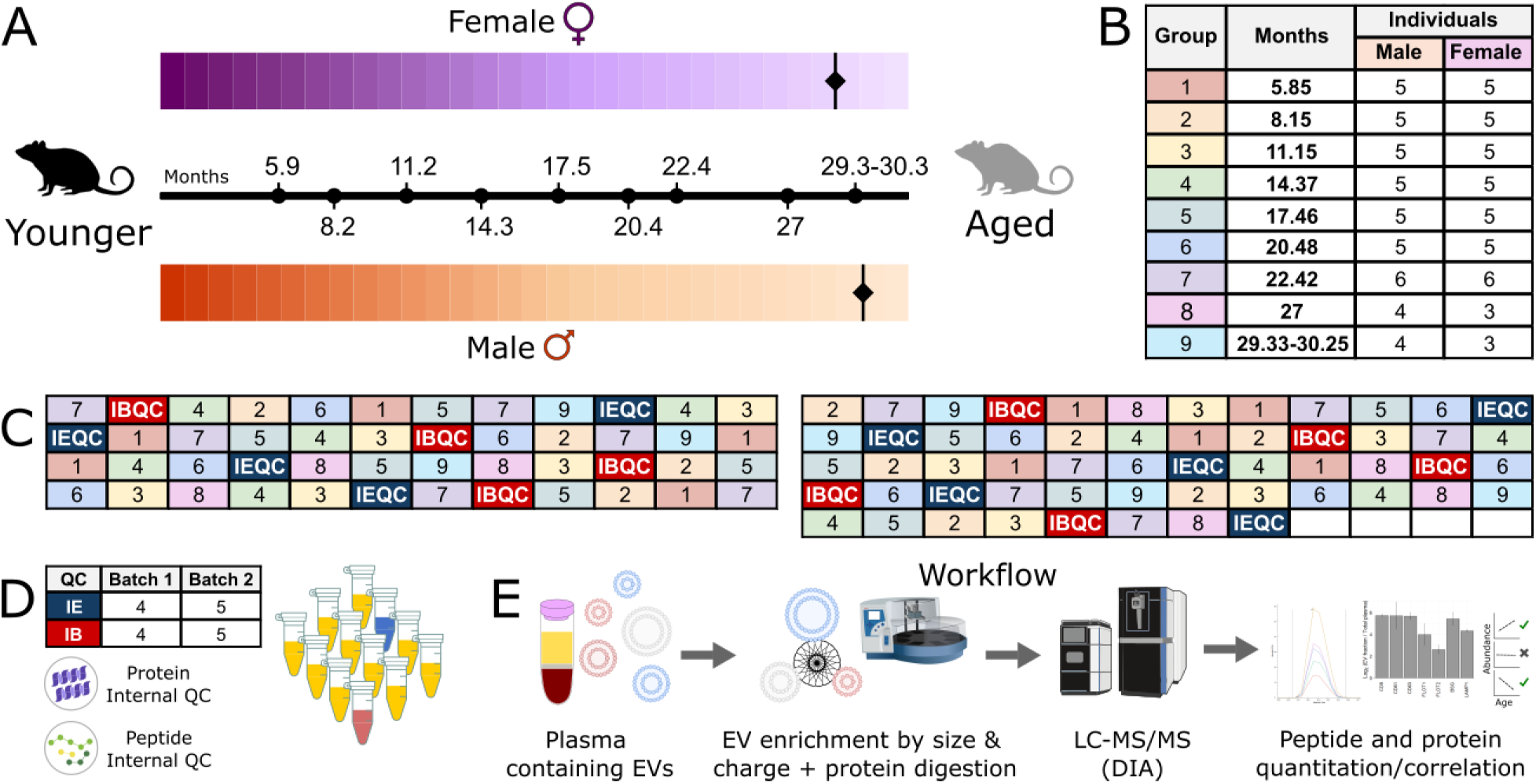
Summary of the aging cohort including age groups for both sexes (A). The median age of male and female C57BL/6J (Yuan et al., 2009) are denoted with a diamond on the timeline of each sex. Experimental sample numbers are listed by age group and sex (B). Plate layouts of the samples across two plates (C) and the number and types of QC samples (D) are listed per batch. Summary of the experimental workflow from EV enrichment through data analysis (E). The Astral MS and Vanquish Neo LC icons were designed by Manon Zuurmond and are used under a CC-BY-4.0 license.

Across the full dataset, 2,575 protein groups were identified from 15,969 peptides. Median normalization improved the CV distribution in both QC sample types (Supplementary Figure 2A-2D) at the peptide level (IB-QC: 28.7% to 24.5%; IE-QC: 33.5% to 28.4%) and protein level (IB-QC: 24.7% to 18.4%; IE-QC: 32.3% to 24.1%). There was no observed effect of acquisition order or sample preparation batch (Supplementary Figure 2E-2H). Hemolysis, assessed qualitatively prior to processing, produced a detectable signal in PCA that was present at both the peptide and protein-level (Supplementary Figure 2I, 2J). The CV of the internal protein QC, yeast enolase, was 20.2% in the raw data and 22.4% after median normalization across all 86 samples, confirming that normalization did not appreciably increase variance in a protein present at a known and consistent quantity.

**Figure 2:**
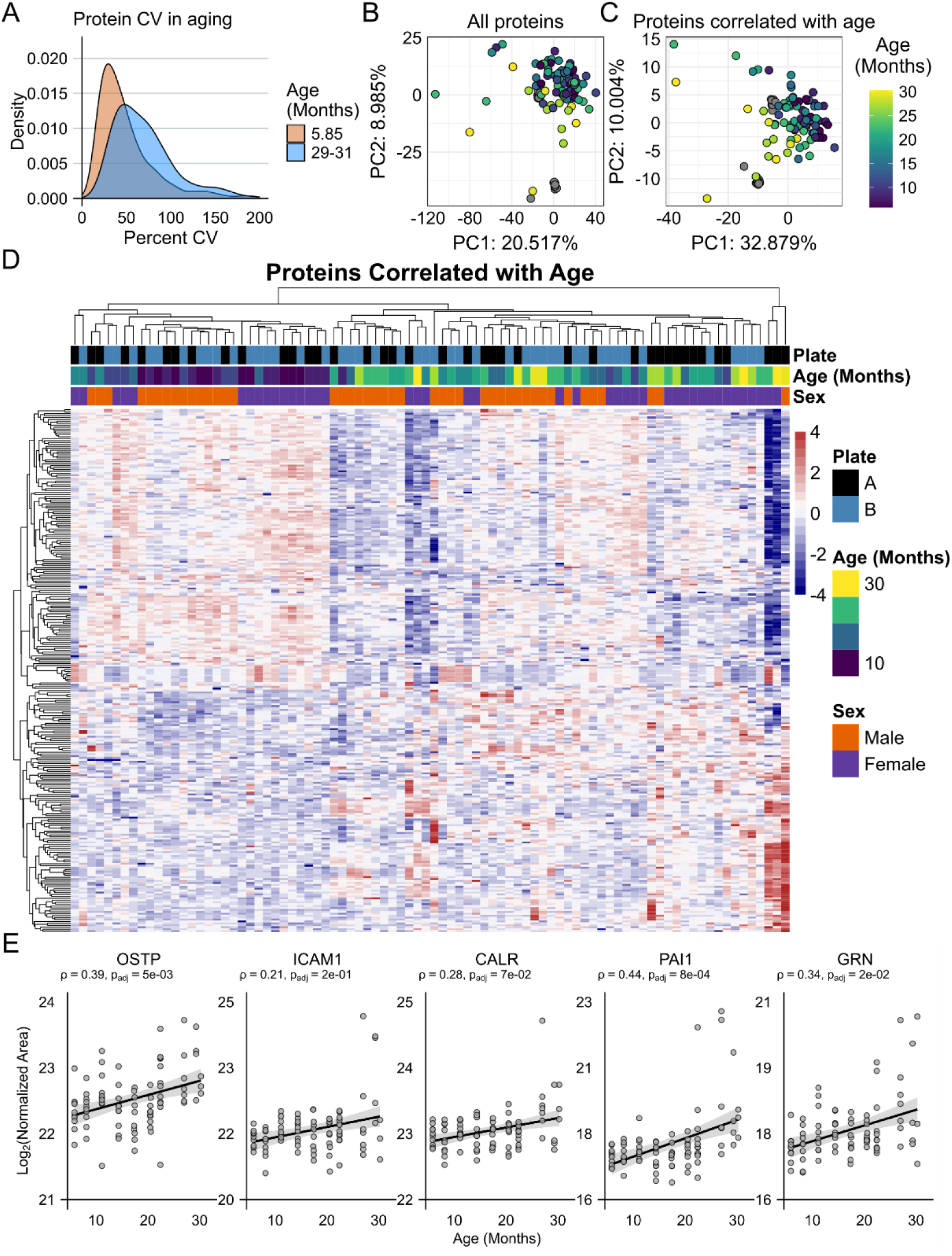
CV density plot of 5.5 (orange) and 29-31 month-old (blue) mice suggests more variability in aged mouse protein abundance (A). The same trend was seen in protein abundance PCA when the points are colored by age (B). After applying Spearman correlation to extract proteins where their abundance increased or decreased with increasing age, the PCA plot reflected a clearer trend associated with age (C). The same proteins correlated with age are plotted in a heatmap with complete clustering (D). Five representative proteins with known associations to senescence, frailty, and aging (OSTP, ICAM1, CALR, PAI1, GRN) are plotted with Spearman Rho and p_adjusted_ (E).

### Protein abundance is more variable in aged mice

Aging is a heterogeneous process that unfolds differently even across genetically identical (inbred) mice. We expected this heterogeneity may be reflected in increased variance in plasma protein abundances as animals age. The median CV in the youngest group of mice (5 months) varied less than the oldest mice (29-31 months) at the protein (Figure 2A) and peptide-level (Supplementary Figure 3A). The PCA plot distribution showed increased spread with increasing age (Figure 2B), with the youngest mice clustering tightly and older mice progressively dispersing which is consistent with growing inter-individual proteome variation across the lifespan.

To determine whether the CVs of protein abundances were significantly different between the youngest and oldest mice, we performed ordinary least squares (OLS) regression. We first compared animals younger than 21 months to those 21 months and older, a threshold corresponding to 70-80% of the median C57BL/6J lifespan and when they are known to be declining in health (Ogiso et al., 2025; Yuan et al., 2009). The β₂ term was positive and significant (Table 1), representing an increased CV of approximately 1.1% per unit on the logarithmic scale of mean abundance in older animals. While modest, this effect is consistent and proteome-wide, likely reflecting a broad erosion of proteome regulation rather than changes in any single protein. Males showed a significantly lower CV than females (β₃ = -0.0072, p < 0.001), a finding explored further in the context of sexually dimorphic aging trajectories below.

**Table 1:** Linear modeling of CV based on mean protein abundance for a given protein in a given age group of mice younger than 21 months or older than 21 months.

| Metric | Coefficient | Standard Error | T | P>\|t\| | [0.025 | 0.975] |
| --- | --- | --- | --- | --- | --- | --- |
| Intercept | 0.1644 | 0.002 | 86.992 | 0.000 | 0.161 | 0.168 |
| Mean Protein Abundance | -0.0090 | 0.000 | -61.609 | 0.000 | -0.009 | -0.009 |
| Age (Cutoff +/- 21 months) | 0.0112 | 0.001 | 15.974 | 0.000 | 0.010 | 0.013 |
| Sex effect (term for Male) | -0.0072 | 0.001 | -10.301 | 0.000 | -0.006 | -0.009 |

We re-applied the model comparing a younger subset of mice (5–15 months) to middle-aged mice (17–21 months) in Supplementary Table 3 and found that while the effect of age was no longer significant, the effect of abundance was essentially unchanged. Together, these results indicate that increased proteome variability is a feature of advanced age specifically, rather than a gradual trend accumulating across the full lifespan.

### Spearman correlation identifies proteins tracking chronological age

Having established that proteome variability increases with age, we next sought to identify which proteins drove this signal and what biological processes they were involved in. Spearman correlation was used to identify peptides and proteins whose abundance significantly correlated with chronological age in male and female mice combined. There were 126 proteins that positively correlated (ρ > 0.3, p_adjusted_ < 0.05) and 146 proteins that negatively correlated (ρ < -0.3, p_adjusted_ < 0.05) with age, yielding 272 proteins derived from 1,536 peptides. When the PCA dataset is narrowed to these 272 proteins, the samples separate well by age (Figure 2C). The normalized abundance of these 272 proteins are plotted on a heatmap with complete clustering (Figure 2D); the clusters separate clearly by age and by sex, providing a visual summary of the coherent age-associated signal in this dataset. The lists of positively or negatively correlated proteins and peptides by Spearman Correlation are available in Supplementary Tables 4 and 5, respectively.

This method identified proteins associated with age that are established in the literature to be involved in the aging process. Five examples are plotted in Figure 2E. Plasminogen activator inhibitor 1 (PAI1), osteopontin (OPN), and Progranulin (GRN) increased with age in both sexes and are associated with aging and senescence (Aversa et al., 2024; Schafer et al., 2020); PAI1 is a key mediator of senescence (Kortlever et al., 2006; Vaughan et al., 2017). Calreticulin (CALR) and GRN have been proposed as biomarkers of frailty in aging (Cardoso et al., 2018). Intercellular Adhesion Molecule 1 (ICAM1) showed a trend consistent with its known involvement in senescence (Aversa et al., 2024; Schafer et al., 2020) but did not reach statistical significance

The five most increased proteins based on Spearman rho are shown in Figure 3A and the five most decreased in Figure 3B. The most increased included Peroxiredoxin-4 (PRDX4, ρ = 0.53), Integral membrane protein 2B (ITM2B, ρ = 0.55), Macrophage colony-stimulating factor 1 receptor (CSF1R, ρ = 0.59), and two cathepsins: B (CATB, ρ = 0.64) and D (CATD, ρ = 0.54). The most decreased were Obscurin-like protein 1 (OBSL1, ρ = -0.67), Lumican (LUM, ρ = -0.69), Leucine-rich repeat neuronal protein 4 (LRRN4, ρ = - 0.67), Fibulin-1 (FBLN1, ρ = -0.66), and Vasorin (VASN, ρ = -0.66).

**Figure 3:**
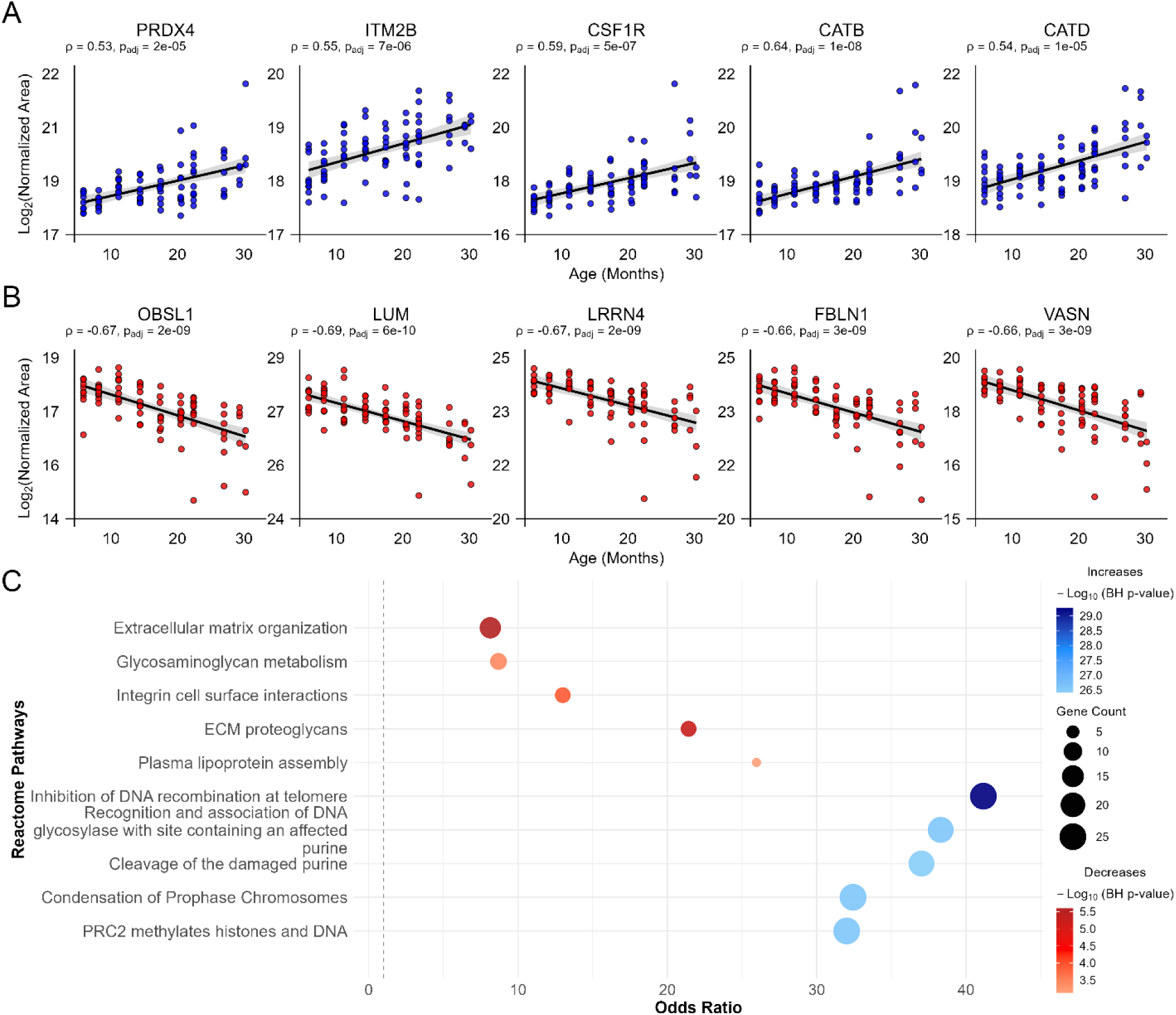
The highest Rho with p_adjusted_ < 0.05 that correlate positively with age (A) and the most negative Rho with p_adjusted_ < 0.05 that correlate negatively with age (B) are shown for all 86 mice in the cohort. Reactome pathway analysis was performed on the proteins that correlated with age across the cohort. The top 5 pathway are shown sorted by the most significant p-value for proteins that increased (blue) or decreased (red) with age (C).

Among the most increased proteins, the thiol-dependent peroxidase PRDX4 regulates H_2_O_2_ levels in the ER and has been linked to reproductive aging and inflammatory NF-κB signaling (Klichko et al., 2016; X. Liang et al., 2020; Tavender & Bulleid, 2010). Notably, ITM2B, CSF1R, CATB, and CATD have each been independently linked to Alzheimer’s or Parkinson’s disease pathology (Del Campo et al., 2014; Dolfe et al., 2018; Hsu et al., 2018; Jiang et al., 2020; Poska et al., 2016; Robak et al., 2017; Rojo et al., 2017), suggesting that the proteins most strongly increasing with age in circulating EVs may reflect early neuroinflammatory or proteostatic changes relevant to neurodegenerative disease.

Among the decreased proteins, OBSL1 is a paralog of the giant muscle scaffold protein obscurin and shares functional redundancy in skeletal and cardiac muscle (Blondelle et al., 2019; Di Paola & Schöck, 2026); its knock-out affects metabolism, mitochondrial size, and autophagy (Fujita et al., 2025), though a direct link to aging has not yet been established. Lumican is a widely expressed small leucine-rich proteoglycan (Nikitovic et al., 2008) that has been considered as a biomarker candidate for open-angle glaucoma (Sharma et al., 2026), vascular aging (Luo et al., 2024), osteoarthritis (Fernández-Puente et al., 2011), and osteosarcopenia (Park et al., 2024), with decreased expression observed in aged murine liver (Liu et al., 2025). LRRN4 is a transmembrane protein involved in hippocampus-dependent learning and long-lasting memory (Bando et al., 2005). FBLN1 and VASN are both part of TGF-β1 signaling, a pro-inflammatory and pro-fibrotic pathway implicated in increasing vascular stiffness; FBLN1 levels have been reduced in aging human aorta (Yasmin et al., 2018), and VASN directly inhibits fibrosis in vascular smooth muscle cells (Pintus et al., 2018; Wang et al., 2020).

### Reactome pathway analysis reveals coherent hallmarks of aging

Using the full list of 272 age-correlated proteins, Reactome pathway enrichment analysis was performed with a significance cutoff of p < 0.05. Of the top 10 pathways positively correlated with age (Figure 3C, blue. Supplementary Table 6), the majority were associated with maintenance of the genome which is a thoroughly established primary hallmark of aging and age-related disease (López-Otín et al., 2023; Niedernhofer et al., 2018; Ribezzo et al., 2016; Schumacher et al., 2021). Of the top 10 pathways negatively correlated with age (Figure 3C, red. Supplementary Table 7), pathways associated with extracellular matrix organization and metabolism stood out, consistent with known ties to the ECM In aging and age-related disease (Haydont et al., 2019; Rahmati et al., 2017; Ramli et al., 2025; Wlaschek et al., 2021) the established antagonistic hallmarks of deregulated nutrient sensing and mitochondrial dysfunction (López-Otín et al., 2023).

Examining the pathways with the largest odds ratios that passed the significance cutoff provides additional resolution. For proteins increasing with age, the emphasis on genomic maintenance pathways remained strong (Supplementary Figure 3B). For proteins decreasing with age, the top hits by odds ratio were dermatan sulfate and chondroitin sulfate biosynthesis and degradation, reinforcing the ECM signal (Supplementary Figure 3C). The next several pathways implicated lipoprotein processing, including plasma lipoprotein assembly, HDL remodeling, and chylomicron assembly and remodeling. These findings connect to highly conserved longevity pathways including mTOR and insulin/IGF1 signaling which are known to intersect with lipid metabolism (Mutlu et al., 2021) and may reflect age-related shifts in circulating lipoprotein biology that are relevant to human cardiometabolic disease (Maranhão et al., 2020).

### Sexual dimorphism in aging plasma EV proteome evaluated by Spearman correlation and OLS interaction

Sexual dimorphism is a well-established phenomenon in mammals that influences biological phenotypes including aging (Hägg & Jylhävä, 2021; Karp et al., 2017). In the full dataset, we found only modest separation between males and females at the peptide (Supplementary Figure 4A) and protein (Supplementary Figure 4B) level, although there was less variance associated with the protein-level data. Once narrowed to the 272 proteins that correlated most with age, separation overall by sex remained limited (Supplementary Figure 4C). This suggests that sex differences in the aging plasma EV proteome are more about trajectory than baseline abundance.

To identify which individual proteins had divergent aging trajectories between sexes, we applied two complementary approaches. First, sex-stratified Spearman correlation of all proteins was run separately in the median-normalized dataset considering only females (n = 42) or only males (n = 44), using the same cutoff of ρ > 0.3 or ρ < -0.3 with p_adjusted_ < 0.05 and FDR thresholds as the full cohort (Supplementary Table 8). Of the five example age-related markers considered in the full cohort analysis, changes with age differed by sex (Supplementary Figure 4D–4H). PAI1, OPN, and GRN increased with age in both sexes. ICAM1 increased in males (ρ = 0.42) and didn’t change (ρ = 0.001) in females. CALR was relatively unchanged across the lifespan in males (ρ = -0.1) and increased in females (ρ = 0.55). These results demonstrate that even well-validated aging markers cannot be assumed to be sex-invariant. The Spearman analysis further yielded 21 proteins with the greatest magnitude of differences between male and female mice based on Spearman rho and calculated FDR, yielding 21 proteins summarized in Table 2. These proteins with a change in rho that passed the FDR filter of 0.05 are summarized in Table 2 and flagged “Yes” the column “Δρ FDR<0.05”. In females, proteins tending to increase with age were associated with coagulation, proteolysis, and protein processing, while those decreasing were associated with the immune system and extracellular matrix. In males, proteins increasing with age were connected to the immune system and ECM processes, while those decreasing broadly covered sex olfactory signaling and proteolysis.

**Figure 4:**
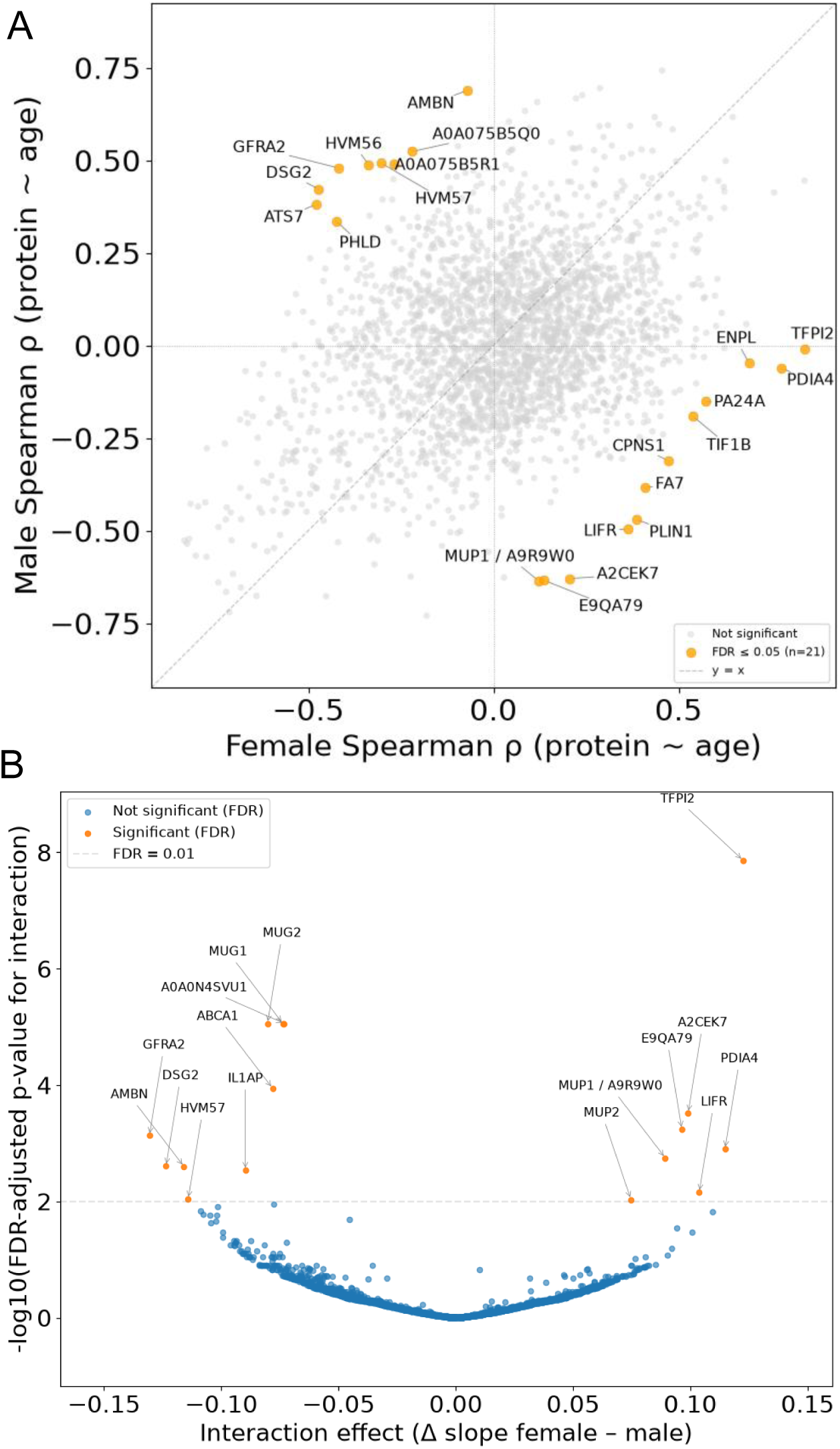
Sex-divergent aging trajectories were identified by complementary statistical approaches. (A) Scatter plot of male versus female Spearman ρ (protein ∼ age) for all detected proteins. Proteins with significantly different age trajectories between sexes after FDR correction (n = 21) are highlighted in orange. (B) Volcano plot of the OLS sex-by-age interaction term (Δ slope, female minus male) versus −log10 FDR-adjusted p-value. Proteins with a significant interaction effect (FDR < 0.01, n = 16) are highlighted in orange. Ten proteins were identified by both approaches.

**Table 2:** The 27 proteins significantly altered with age between the sexes either by significant Spearman Rho differences or by significant OLS interaction terms. Proteins with a significantly different change in spearman Rho and FDR cutoff below 0.05 are flagged “yes” in the column “Δρ FDR<0.05”. The male (ρ M) and female (ρ F) spearman rho and FDR of the change in rho (Δ ρ FDR) are included. The proteins with a significant age and sex interaction by OLS Are flagged “yes” in the column “OLS Int β q<0.01”. The interaction terms (Int β) and q value (Int q) are also included. Broad, manually-defined categories were identified and listed under “Functional Categories”. 10 proteins overlapped between the Spearman and OLS analysis and are flagged “yes” in both columns.

| Protein | Accession | $\Delta \rho$<br>FDR<0.05 | $\rho$ M | $\rho$ F | $\Delta \rho$ FDR | OLS Int $\beta$<br>q<0.01 | Int $\beta$ | Int q | Functional Categories |
| --- | --- | --- | --- | --- | --- | --- | --- | --- | --- |
| TFPI2 | O35536 | Yes | -0.01 | 0.84 | 9.60E-05 | Yes | 0.122 | 1.40E-08 | Coagulation \| proteolysis |
| FA7 | P70375 | Yes | -0.38 | 0.41 | 3.10E-02 | No | 0.075 | 1.30E-01 | Coagulation \| proteolysis |
| DSG2 | O55111 | Yes | 0.42 | -0.47 | 1.10E-02 | Yes | -0.124 | 2.50E-03 | Extracellular matrix |
| ATS7 | Q68SA9 | Yes | 0.38 | -0.48 | 1.20E-02 | No | -0.102 | 2.20E-02 | Extracellular matrix |
| AMBN | O55189 | Yes | 0.69 | -0.07 | 1.20E-02 | Yes | -0.116 | 2.50E-03 | Extracellular matrix |
| MUG1 | P28665 | No | -0.53 | -0.83 | 1.90E-01 | Yes | -0.074 | 8.80E-06 | Immune response \| protease inhibitor |
| LIFR | P42703 | Yes | -0.49 | 0.36 | 1.20E-02 | Yes | 0.104 | 7.00E-03 | Inflammation or Immune response |
| HVM56 | P18527 | Yes | 0.49 | -0.34 | 1.60E-02 | No | -0.109 | 1.50E-02 | Inflammation or Immune response |
| HVM57 | P18528 | Yes | 0.49 | -0.3 | 2.40E-02 | Yes | -0.114 | 9.10E-03 | Inflammation or Immune response |
| IGHV5-17 | A0A075B5R1 | Yes | 0.49 | -0.27 | 4.30E-02 | No | -0.092 | 7.20E-02 | Inflammation or Immune response |
| PA24A | P47713 | Yes | -0.15 | 0.57 | 4.30E-02 | No | 0.092 | 6.50E-02 | Inflammation or Immune response |
| IGHV5-6 | A0A075B5Q0 | Yes | 0.52 | -0.22 | 4.30E-02 | No | -0.095 | 5.50E-02 | Inflammation or Immune response |
| IL1AP | Q61730 | No | -0.08 | -0.63 | 1.40E-01 | Yes | -0.09 | 2.90E-03 | Inflammation or Immune response |
| PLIN1 | Q8CGN5 | Yes | -0.47 | 0.39 | 1.20E-02 | No | 0.109 | 1.50E-02 | Lipids and Lipoproteins |
| PHLD | O70362 | Yes | 0.34 | -0.42 | 4.30E-02 | No | -0.105 | 1.70E-02 | Lipids and Lipoproteins |
| ABCA1 | P41233 | No | -0.43 | -0.8 | 1.90E-01 | Yes | -0.078 | 1.10E-04 | Lipids and Lipoproteins |
| GFRA2 | O08842 | Yes | 0.48 | -0.42 | 1.10E-02 | Yes | -0.131 | 7.40E-04 | Neuronal survival \| kinase signalling |
| TIF1B | Q62318 | Yes | -0.19 | 0.54 | 4.90E-02 | No | 0.101 | 3.30E-02 | Nuclear protein |
| MUP12 | A2CEK7 | Yes | -0.63 | 0.21 | 1.10E-02 | Yes | 0.099 | 3.00E-04 | Olfactory signal |
| MUP7 | E9QA79 | Yes | -0.63 | 0.14 | 1.60E-02 | Yes | 0.096 | 5.80E-04 | Olfactory signal |
| MUP1 | P11588 | Yes | -0.64 | 0.12 | 1.90E-02 | Yes | 0.089 | 1.80E-03 | Olfactory signal |
| MUP2 | P11589 | No | -0.63 | 0.03 | 6.80E-02 | Yes | 0.075 | 9.30E-03 | Olfactory signal |
| MUG2 | P28666 | No | -0.43 | -0.82 | 1.20E-01 | Yes | -0.08 | 8.80E-06 | Protease inhibitor |
| MUG5 | A0A0N4SVU1 | No | -0.54 | -0.83 | 2.20E-01 | Yes | -0.073 | 8.80E-06 | Protease inhibitor |
| PDIA4 | P08003 | Yes | -0.06 | 0.78 | 1.10E-03 | Yes | 0.115 | 1.30E-03 | Protein processing |
| ENPL | P08113 | Yes | -0.05 | 0.69 | 1.60E-02 | No | 0.094 | 2.80E-02 | Protein processing |
| CPNS1 | O88456 | Yes | -0.31 | 0.47 | 3.20E-02 | No | 0.09 | 8.30E-02 | Proteolysis |

The second approach we used was OLS analysis to identify sexually divergent proteins in this dataset considering a sex-specific age effect at the protein level. This analysis yielded 16 proteins with a significant q-value (q < 0.01) after FDR correction (Figure 4, Supplementary Table 9). These included 10 of the 21 proteins identified by the sex-stratified Spearman analysis, providing convergent support for the most robust findings. The OLS results for these significant 16 proteins are summarized in Table 2 and flagged “Yes” the column “OLS Int β q<0.01”. The interaction effect was higher in females for Tissue factor pathway inhibitor 2 (TFPI2), MUP12, PDIA4, MUP1, and LIFR; in males, the interaction effect was higher for GFRA2, DSG2, AMBN, and HVM57. Six proteins were captured exclusively by OLS including MUG1, MUG2, MUG5, MUP2, ABCA1, and IL1AP, broadly related to immune function, protease inhibition, and the ECM remodeling.

### Sex-divergent biological themes in aging

Taken together, the proteins with the most sex-divergent aging trajectories by sex-stratified Spearman analysis and OLS fell into two broad biological themes.

In females, the most prominent signal was an increase in proteins associated with coagulation and proteolysis. Most striking was TFPI2 (ρ♀ = 0.840), the most strongly age-correlated protein in females by either analysis that has been linked to coagulation, fibrinolysis, fertility, and potentially cancer (Altmäe et al., 2011; Ito et al., 2024; Kobayashi et al., 2023; Ota et al., 2021; Wojtukiewicz et al., 2024). The most decreased protein in females was ATS7 (ADAMTS-7, ρ♀ = -0.479), which has been implicated in osteoarthritis, collagen-induced arthritis, and atherosclerosis (Bengtsson et al., 2017; Lai et al., 2014; Zhang et al., 2015). ATS7 increased in males with age (ρ♂ = 0.381), representing one of the clearest sex-divergent trajectories in the dataset.

In males, the most striking age-related decreases were in three MUP proteins: MUP1 (ρ♂ = -0.635), MUP7 (ρ♂ = -0.634), and MUP12 (ρ♂ = -0.629), all known to be more highly expressed in male mice. MUP proteins are known to decline in senescent male mouse urine (Garratt et al., 2011), and we speculate this could reflect a reduced energetic investment in olfactory signaling focused on reproduction, which would be consistent with the disposable soma theory of aging (Kirkwood, 1977). Ameloblastin (AMBN, ρ♂ = 0.689) was the most increased protein in males which was an unexpected finding given its known role in tooth enamel mineralization (Kegulian et al., 2024). However, AMBN contains an alpha-helix that embeds within synthetic membranes and may be transported through them (Su et al., 2019), and oral health declines in aging mice whose teeth grow throughout their lifespan (Catón & Tucker, 2009; S. Liang et al., 2010)

We observed other proteins with established sexually dimorphic expression whose age trajectories diverged by sex, including the MUG proteins (MUG1, MUG2, MUG5), which differ between males and females in mouse circulation (Han et al., 2022, 2022) and showed a stronger interaction effect in males by OLS. Collagen alpha-1(VI) chain (CO6A1) declined in females (ρ♀ = -0.544) but remained steady in males (ρ♂ = -0.096); CO6A1 knock-out mice have been considered a model of muscle aging (Capitanio et al., 2017) and sarcopenia is more common in female mice (Kerr et al., 2024) and humans (Geraci et al., 2021; Yang et al., 2019).

### Circulating plasma aging biomarkers predict chronological age

It is unlikely any single protein will be sufficient to serve as a biomarker of the aging process, which is inherently heterogeneous. While individual proteins may differ in terms of direction and magnitude of correlation with age, we sought to evaluate whether the combined contribution of the ∼2,500 proteins identified across this cohort could be leveraged to identify a multivariable signature of aging capable of predicting chronological age. We used Elastic Net regularized linear regression (Zou Hastie 2005 J Roy Stat Soc Ser B) to determine whether proteins measured in this cohort of aged mice could be used to predict chronological age, and which proteins contributed most to that prediction. Performance was estimated using 10-fold cross-validation repeated 5 times. The estimated MAE was 2.75 months and the estimated r² was 0.79 (Figure 5A). In the final model, 277 proteins out of 2,575 proteins measured contributed a non-zero coefficient (Supplementary Table 10).

**Figure 5:**
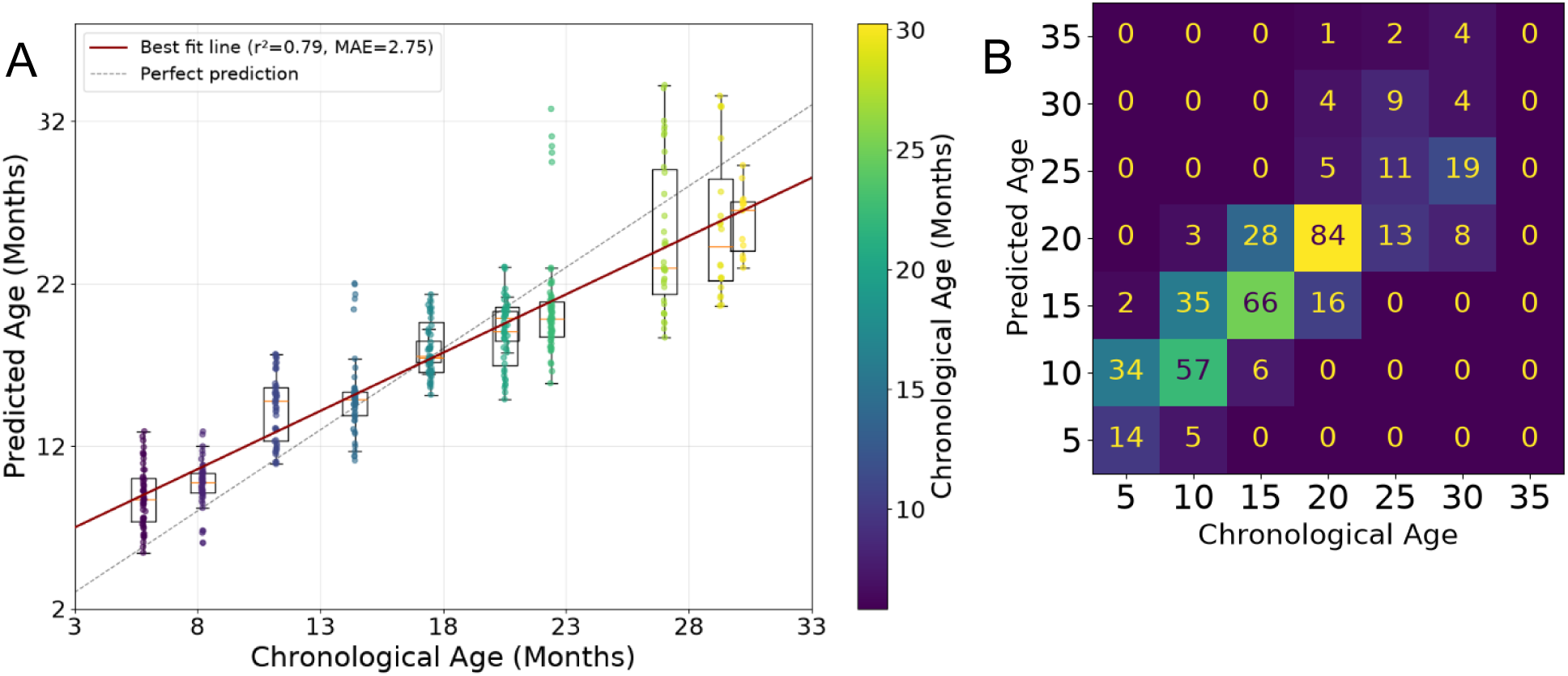
Plot of the 10-fold cross-validation with 5 repeats of the Elastic Net model to predict chronological age using the median-normalized, logarithmically-transformed proteins abundances measured in the full dataset of 2,575 proteins (A). The confusion matrix of this model (B) illustrating that the model performs best in the middle of the age-range tested and is less accurate at predicting chronological age at the earliest and latest ages of the cohort.

The confusion matrix indicates that prediction of chronological age from protein abundance becomes less accurate at the extremes of the age range in either the youngest and oldest mice (Figure 5B). The reduced accuracy in the age prediction with increasing age is consistent with the increased variability in protein abundance we observed as the mice aged (Figure 2A, B, and C). A final model was constructed using all data and the protein coefficients from this model are reported in Supplementary Table 11.

Of the 277 proteins in the model, 113 overlapped with the significant Spearman-correlated proteins across the full cohort (Supplementary Table 12) and 178 considering the sex-differential Spearman analysis (Supplementary Table 13).

Of the 15 proteins with the most negative or the 15 most positive coefficients in the Elastic Net model, 9 were proteins identified within the 15 most decreased or increased with age by Spearman correlation in either the full dataset or in either males or females, indicating broad concordance between the two analytical approaches.

We applied the same Elastic Net approach to the 15,969 peptides identified in the dataset to determine whether peptide-level data captured aging signal beyond what was accessible at the protein level. The estimated peptide MAE was 3.03 and r² was 0.75 (Supplementary Figure 5A, 5B), suggesting a modestly greater disparity between predicted and actual chronological age relative to the protein-level model (Supplementary Table 14, 15). We postulated that protein-level data which was generated by summing peptide intensities, may reduce the impact of natural variation associated with individual proteoforms and post-translational modifications that are likely to be heterogeneous in aging. Using OLS analyses, we identified 96 peptides with significant age-associated q-values at the peptide level that originated from proteins that did not show a significant association with age in the corresponding protein-level analysis (Supplementary Figure 5C, Supplementary Table 16). These peptides may reflect age-related changes that would not be captured in protein-level analyses alone. The most significant peptides represented proteins involved in cytoskeletal rearrangement (FLNA, ADDA), clotting (FA10, FA9), and oxidative stress (PRDX1). When applying a q-value significance threshold of 0.01, no cases were observed in which peptide- and protein-level age coefficients were in opposite directions; effect estimates were directionally concordant between peptide and protein levels in all cases where both reached significance. However, the peptide-level clock still performed well (r² = 0.75), and its power may be improved by evaluating peptide-level data in a larger validation cohort or across multiple species.

## Discussion

Here we show that a species-agnostic, mass spectrometry-based approach to EV-enriched plasma proteomics can identify biologically coherent signatures of aging, generate a predictive proteomic clock, and reveal sex-specific aging trajectories. By coupling Mag-Net EV enrichment (Wu et al., 2025) to DIA-MS, this approach circumvents the cross-species translation failures and ambiguous signal attribution that limit affinity-based platforms, while achieving proteome depths that reveal both established aging hallmarks and novel candidate markers. These findings suggest that proteomic clocks derived from circulating EVs represent a complementary approach to epigenetic clocks (Horvath & Raj, 2018) grounded in the functional output of the genome and epigenome that is accessible from a minimally invasive biofluid.

Heterogeneity is an established hallmark of aging that has been observed in humans, genetically homogeneous model organisms, and within individual tissues (Carnes & Olshansky, 2001; Herndon et al., 2002; Martin, 2012; Mendenhall et al., 2021; Mitnitski et al., 2017). We similarly found that the CV of protein abundance increased with advancing age (Figure 2A) and could be predicted using linear modeling of protein abundance considering age and sex (Table 1, Supplementary Table 3). This increased variation may reflect erosion of epigenome and genomic stability that would impact proteome-wide regulation. This change that was detectable in a small volume of plasma may serve as a metric of biological aging distinct from any individual protein marker.

The proteins most strongly correlated with age in the circulating EV proteome recapitulate established aging hallmarks (López-Otín et al., 2023). Proteins increasing with age were enriched in genome maintenance pathways which represent a primary hallmark of aging and are implicated in age-related diseases including cancer (Niedernhofer et al., 2018; Ribezzo et al., 2016; Schumacher et al., 2021).The decreasing proteins implicated ECM organization, lipoprotein metabolism, and lipid processing pathways connected to conserved longevity signaling including mTOR and insulin/IGF1 (Haydont et al., 2019; Lacolley et al., 2017; Mutlu et al., 2021; Rahmati et al., 2017; Ramli et al., 2025; Wlaschek et al., 2021). Among the known senescence and frailty markers recovered, PAI1, osteopontin, and progranulin increased in both sexes, while calreticulin showed a female-specific increase and ICAM1 showed divergent trajectories by sex, underscoring that even well-validated aging markers cannot be assumed to be sex-invariant.

Among the novel candidates, we noted a striking convergence of the proteins most strongly increasing with age on known pathology of Alzheimer’s and Parkinson’s disease. ITM2B, CATB, CATD, and CSF1R have each been independently linked to neuroinflammation, amyloid processing, or microglial function (Hsu et al., 2018; Poska et al., 2016; Robak et al., 2017; Rojo et al., 2017). Recent reviews have described how circulating EVs may carry cargo from the central nervous system and connect to neurodegenerative diseases (Dutta et al., 2023; Hornung et al., 2020). Whether our results reflect neuroinflammatory cargo being transmitted throughout the body via EVs prior to symptom onset or whether these increases reflect systemic aging processes that parallel central nervous system changes remains an open question. Nonetheless, this convergence motivates prospective investigations of plasma EV proteomics as a tool for early neurodegenerative disease surveillance.

The sexually dimorphic aging trajectories identified here reflect distinct biological themes in males and females. In females, the dominant signal was an increase in coagulation and proteolysis-related proteins with age, most prominently TFPI2, alongside decreases in immune and ECM proteins. In males, the most striking age-related changes were decreases in MUP proteins which are consistent with reduced energetic investment in reproductive signaling under the disposable soma framework (Kirkwood, 1977) and increases in immune and ECM-related proteins. Broadly, females appear to age at the proteome level in a manner reflecting shifts in hemostasis and proteolysis, while males exhibit a more prominent immunosenescence signature. These patterns are consistent with known sex differences in aging trajectories and age-related disease prevalence (Hägg & Jylhävä, 2021) and may help explain why certain age-related conditions differ in onset and severity between the sexes.

The Elastic Net proteomic clock achieved an MAE of 2.75 months and r² of 0.79 at the protein level, with a comparable peptide-level model (MAE = 3.03, r² = 0.75). Reduced accuracy at the youngest and oldest ages likely reflects, respectively, compressed proteome differences early in life and the increased inter-individual heterogeneity that characterizes advanced age. This finding connects clock performance directly back to the variability findings. The identification of 96 peptides with significant age associations whose parent proteins did not reach significance at the protein level suggests that proteoform- and PTM-level changes represent an additional layer of aging biology not captured by protein-level aggregation alone.

Relative to clocks generated using large-scale human data from thousands of patients tracked in the UK Biobank derived from plasma aptamer platform data, we observed a lower r² value in our clock and a larger MAE. Using a human dataset collected from 491 SOMAmers, the best clock identified by Lehallier et al. observed a Pearson correlation of 0.96 and a MAE of 2.44 years in their test set data (Lehallier et al., 2020). Relative to the average of the male and female median lifespan of C57BL/6J (Yuan et al., 2009), the MAE of 2.75 months was representative of 9.5% of the median lifespan. In the Lehallier study, their MAE of 2.44 years in the human cohort in the UK Biobank represented 3.4% of the mean age at death of 72.3 years based on the September 24, 2025 report (UK Biobank, 2025). As part of an organ-specific clock study, Oh et al. used LASSO to produce a conventional aging model clock in plasma from 4,778 proteins trained in 1,398 humans with a Pearson correlation of 0.92 (Oh et al., 2023). Our study cohort was far more limited in total individuals evaluated (86 mice) but showed promise for subsequent validation in additional species and larger mouse cohorts.

This study has several important limitations. The cross-sectional design precludes distinguishing biological from chronological aging at the individual level, and longitudinal validation will be essential to determine whether the proteomic clock tracks interventions or health trajectories. The cohort was limited to a single inbred strain (C57BL/6J), and findings may not generalize across genetic backgrounds. Sample sizes per age and sex group are modest (3–6 animals), limiting statistical power for interaction analyses. No orthogonal validation of individual protein candidates was performed, and replication in an independent cohort is needed before any marker is advanced as a biomarker candidate.

The findings presented here lay groundwork for several important directions. With the recent explosion of studies seeking biomarkers and signatures of aging, consideration into the initial evaluation, validation, and transparent publication of study findings has been undertaken. Recommendations to generate reliable aging biomarkers have been well-reviewed elsewhere (Moqri et al., 2024; Tanaka et al., 2025) and will be crucial to ensure that work laboratory geroscience can be translated to the clinic in the future. We aim to consider non-linear, logarithmic, and biphasic trajectories in future analyses of this dataset.

Of the protein markers most suited for subsequent orthogonal validation as novel aging biomarkers, we propose focusing on those most correlated across the entire cohort and those most sexually divergent. Across the cohort, CSF1R, ITM2B, CATB and CATD increased the most, LRRN4 decreased, and all have ties to neurodegeneration and could be vetted in subsequent studies. In female mice, the clearest increase in TFPI2 that was unchanged in males is noteworthy for follow-up. TFPI2 serum levels measured in 241 human donors (Ito et al., 2024) were not significantly different on average or across ages by Spearman correlation between males and females. However, platelets are known to carry TFPI2 (Vadivel et al., 2014) but are depleted from serum. Because we isolated EVs from plasma, our dataset would include EVs derived from platelets that also circulate in the blood. This may explain the sex-divergent aging trajectory of TFPI2 observed here and warrants further investigation. Other groups have begun to undertake longitudinal analyses (Shen et al., 2024). The inclusion of additional physiological metrics (Dey et al., 2025) may further clarify the mechanisms driving proteome-wide changes with aging and lead to more robust signatures of aging that could be correlated with healthspan. Finally, consistent with recommendations from the field (Moqri et al., 2024; Tanaka et al., 2025), cross-species replication will be an important consideration for aging-specific biomarker studies. Truly conserved processes representative of the aging process would be expected to replicate in disparate species with differing lifespan, environment, and age-related comorbidities. The species-agnostic nature of Mag-Net makes this a tractable next step, and a follow-up cross-sectional study in companion dogs is in progress.

## Acknowledgements

The authors wish to thank all members of the MacCoss laboratory, Brook Nunn and her laboratory, Mariya Sweetwyne, Peter Rabinovitch, Matt Kaeberlein, Daniel Promislow, Jessica Young, Jonathan An, Devin Schweppe, and Katarina Vlajic for their thoughtful discussions about this work as it progressed. We also thank the teams that cared for the mice that made this work possible.

## Author Contributions

Study conception: K.A.T., C.C.W., G.A.C., and M.J.M.

Methodological design included: K.A.T., G.E.M., C.C.W., G.R.K., L.R., A.L.

Experimental work: K.A.T., G.E.M., G.R.K., R.S.J.

Data analysis: K.A.T., M.R., A.M., R.S.J. Study supervision: G.A.C., M.J.M.

Initial manuscript draft: K.A.T., M.R., M.J.M.

Manuscript revisions: K.A.T., M.R., R.S.J., A.L., L.R., G.A.C.

All authors reviewed and approved the final manuscript.

## Data Availability Statement

All raw files, Skyline documents, processed results used as input for figure generation, FASTA files, EncyclopeDIA files, metadata, Nextflow workflow configuration files and output are available on PanoramaWeb ([Murine Plasma EVs](https://panoramaweb.org/MacCoss/Collaborations/Shock-Jax-Churchill/Plasma%20EVs/project-begin.view)). The data input <25 MB, R Markdown files, and Jupyter Notebooks used to generate Figures 2-5, Supplementary Figures 1-5, and Supplementary Tables 1-15 are available on GitHub: (manuscript-aging-mouse-ev). Any input files larger than 25 MB are noted in the ReadMe and freely available on PanoramaWeb.

