## Supplementary material for "Circulating extracellular vesicles in plasma carry accessible molecular signatures of aging in mice": 20260727_supplemental_material

**Suplementary Material for Manuscript**

### Title

### Supplementary Figures

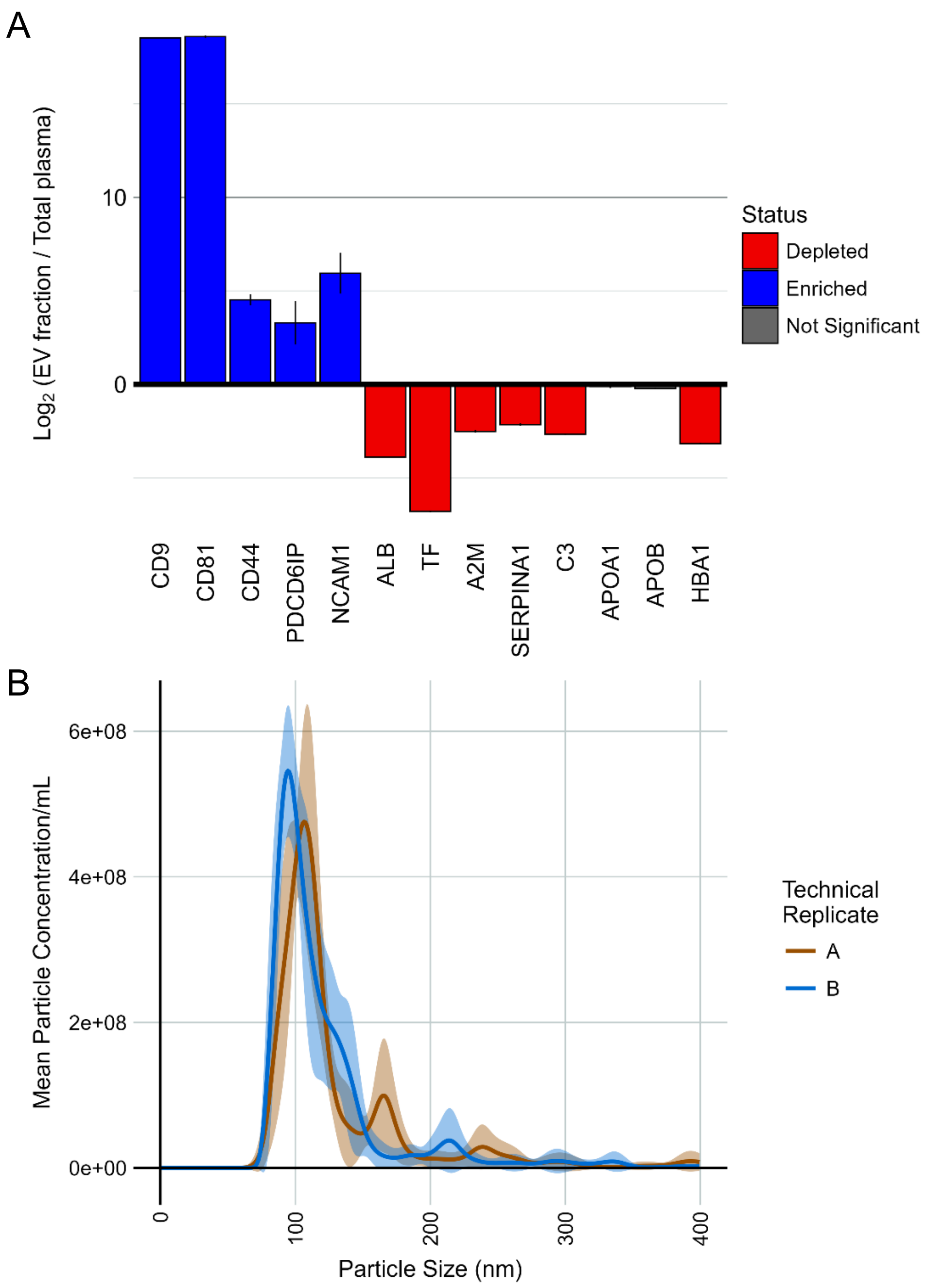

**Supplementary Figure 1:** We confirmed that like previous studies in human plasma the Mag-Net protocol shows an enrichment of established EV markers (A) and a depletion of many high abundance proteins (B) that comprise the majority of the plasma proteome's mass. Nanosight particle analysis also confirmed the presence of particles isolated using Mag-Net (C).

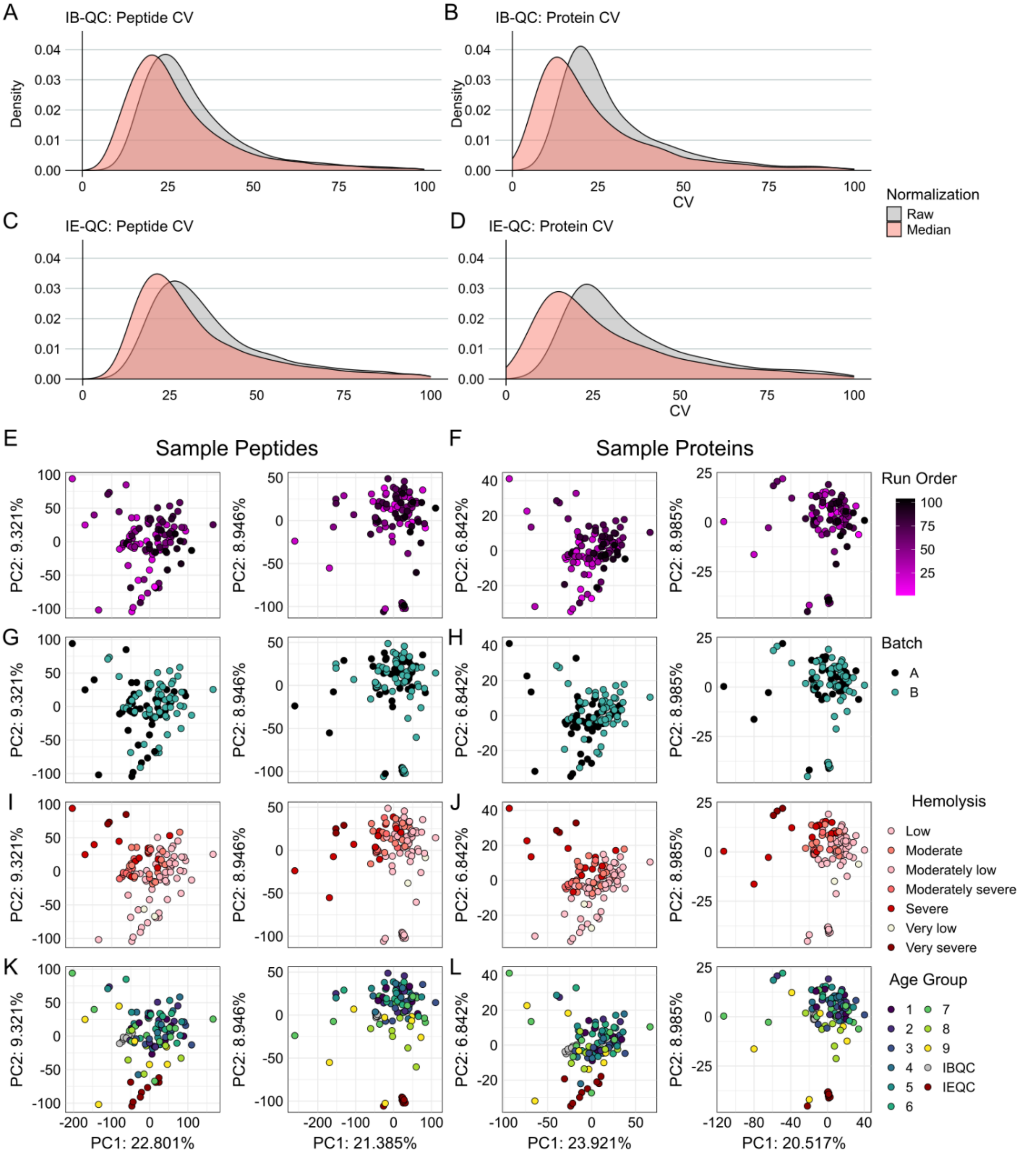

**Supplementary Figure 2:** The distribution of CVs of the peptides and proteins detected in the raw and median-normalized data were evaluated in the IB-QC and IE-QC. Median normalization shifted the distribution of CVs lower in the peptides (A) and proteins (B) of the IB-QC sample and in the peptides (C) and proteins (D) of the IE-QC sample. PCA plots colored by different variables highlighted a few points. First, we found no evidence of a run order effect at the peptide (E) or protein-level (F) with or without median normalization. There was not clear evidence of a sample preparation batch effect with or without median normalization at the peptide (G) or protein (H) level. Some hemolyzed samples were present, and could be observed in at the peptide (I) and protein (J) level. The IB-QC was more hemolyzed than the IE-QC, and this was visualized in a cluster in (I) and (J). Finally, when colored by QC type and by the age groups (bins) described in Figure 1, we observed clear clustering of each control together in the raw data that became tighter after median normalization at the peptide (K) and protein (L) level. We also saw increasing spread of the data with increasing age. The youngest mice in the lowest age group numbers tended to be more tightly clustered than the oldest mice in the highest age group numbers. This increased spread with age was better visualized after median normalization in peptides (K) and proteins (L).

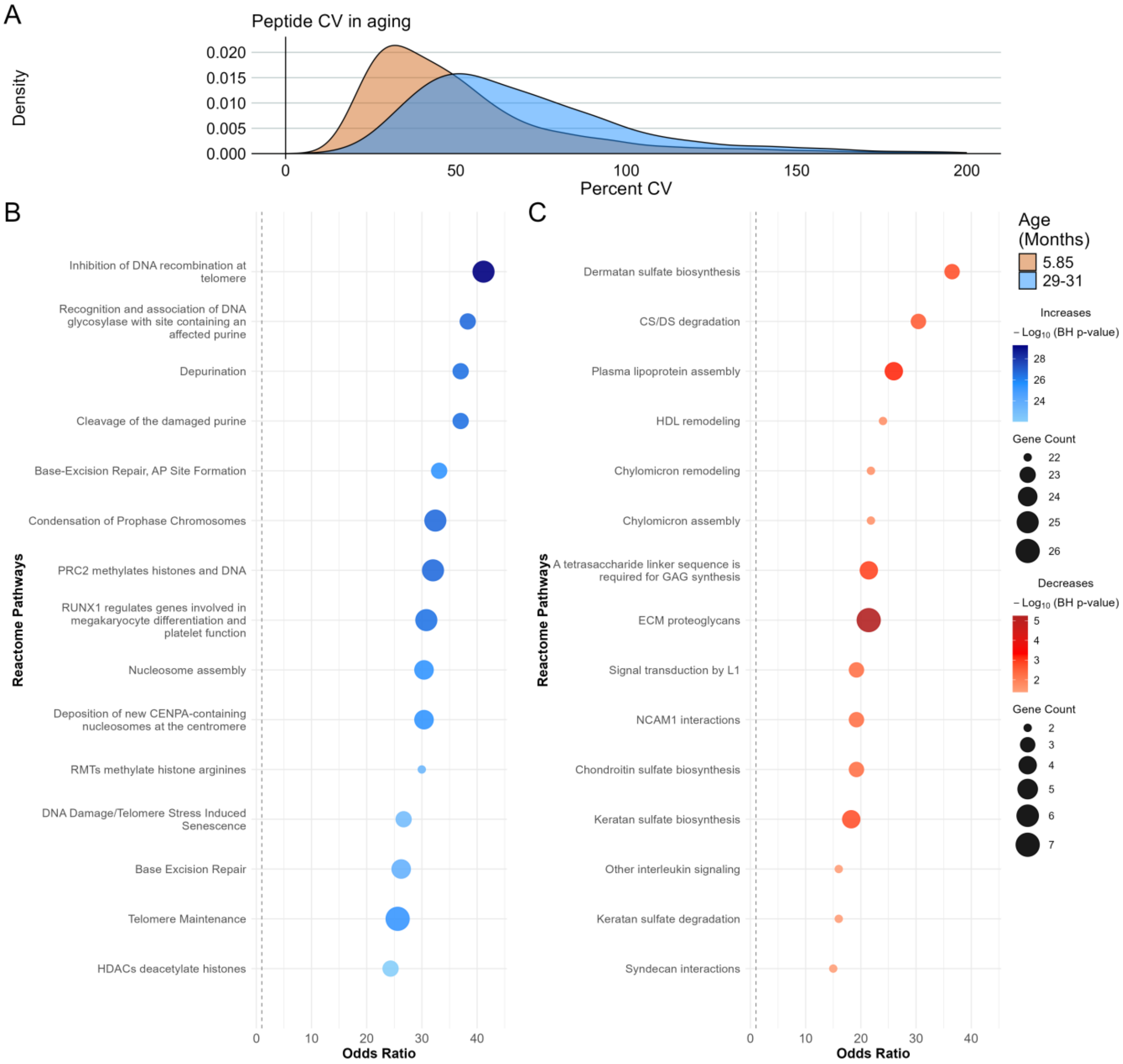

**Supplementary Figure 3:** Density plot of the peptide CV for mice in the 5.5 month (orange) and 31 month (blue) age group indicates that there is more variability in aged mouse peptide abundance (A). Spearman correlation was applied to extract proteins where their abundance increased or decreased with increasing age, and Reactome pathways associated with proteins that correlated positively (B) and negatively (C) were evaluated using the ReactomePA R package. The top 15 pathways with BH-corrected p-value $< 0.05$ and the highest Odds Ratios are plotted.

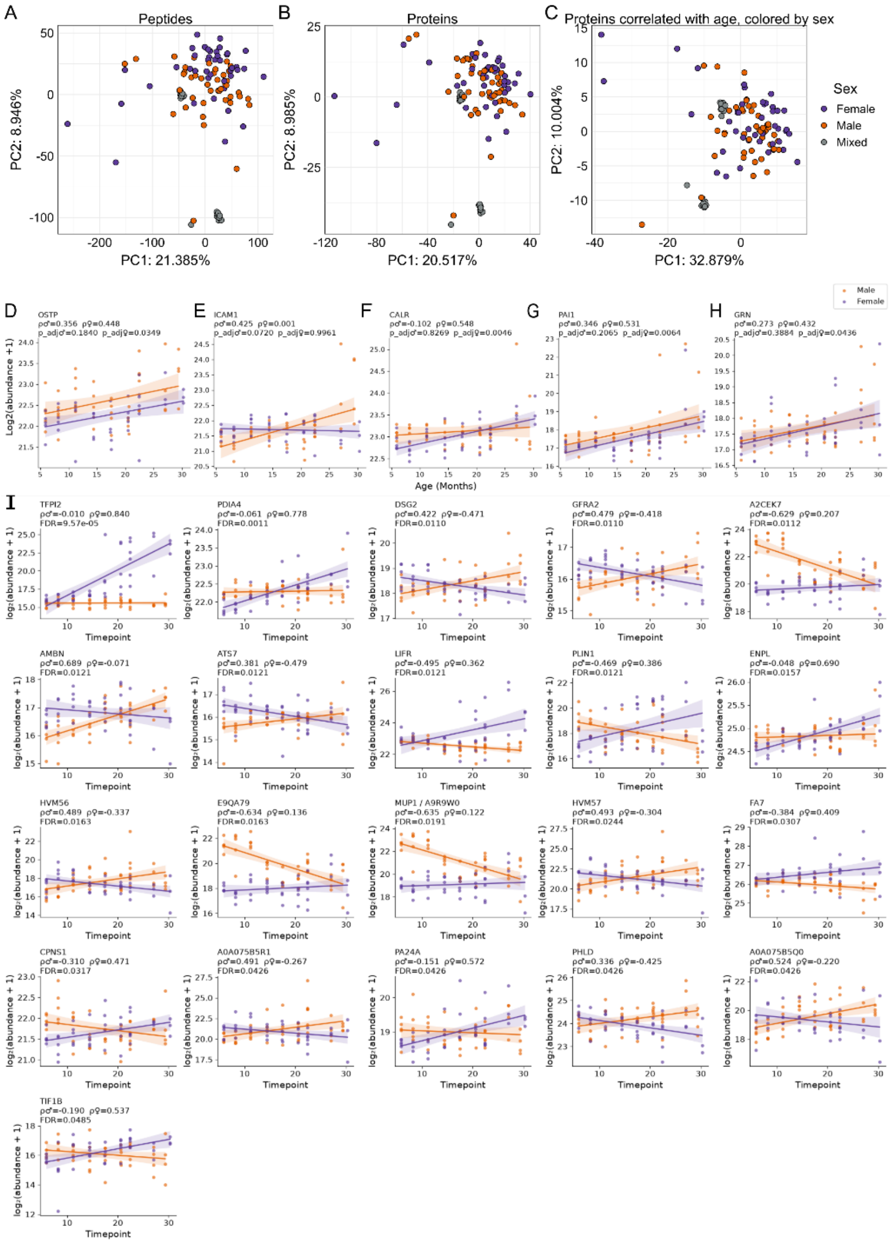

**Supplementary Figure 4:** PCA at the peptide level (A). PCA at the protein level (B). PCA at the protein level filtering the dataset for the proteins with the Spearman Rho greater than 0.3 or less than -0.3 with a p-adjusted < 0.05 after Benjamini-Hochberg correction (C). After re-calculating Spearman correlation at the protein-level for each sex separately, the same known age and senescence-related markers highlighted in Figure 2E are plotted in (D), (E), (F), (G), and (H) split by sex. There are 21 proteins with p < 0.05 after multiple hypothesis testing correction that have the largest difference in Spearman Rho with increasing age between males and females (I). Males and females are plotted in orange and purple, respectively. We used OLS to examine what proteins had a sex-specific age and found 16 had a significant q-value after FDR correction. This included 10 proteins that overlapped with the Spearman analyses.

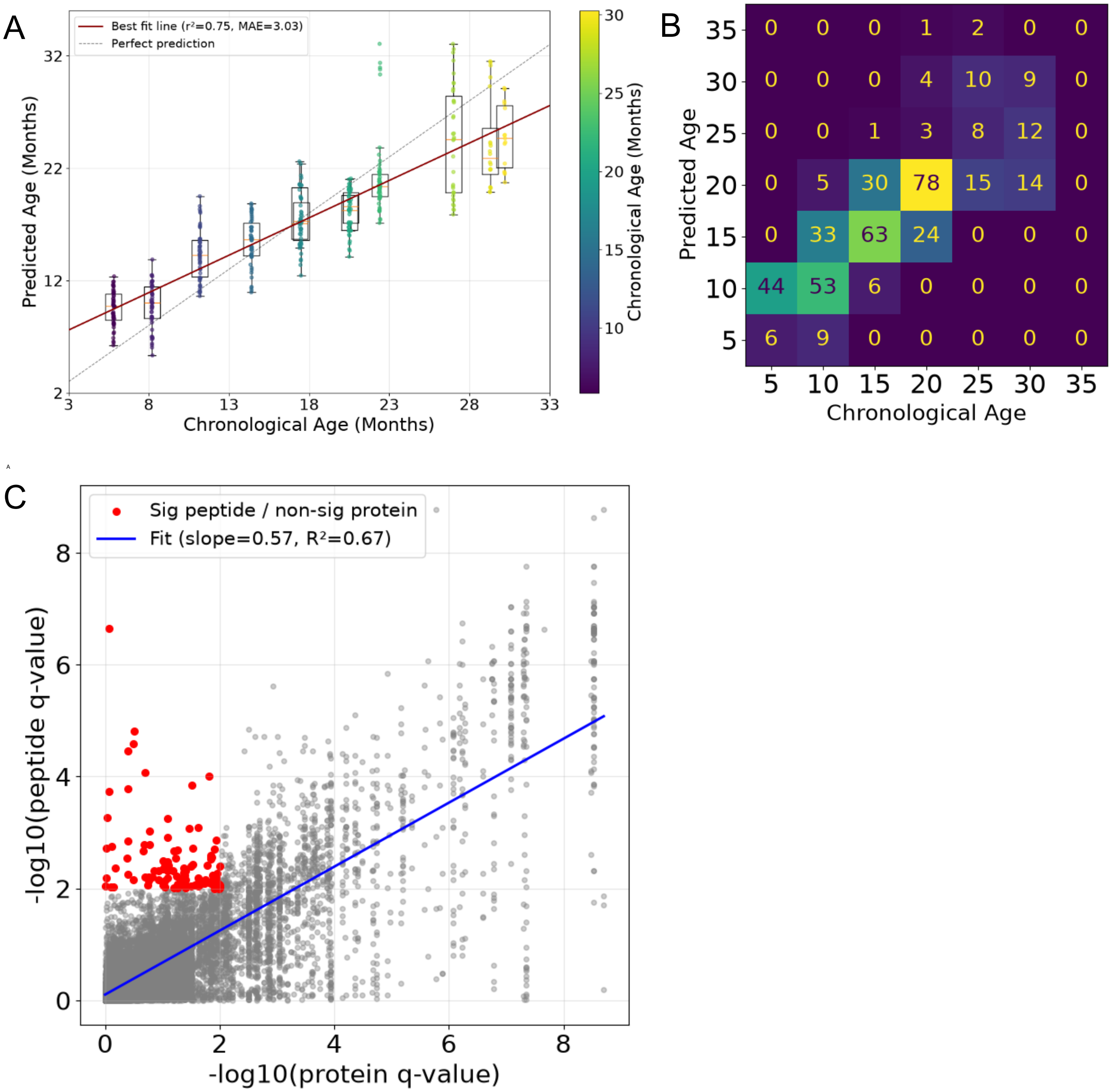

**Supplementary Figure 5:** Plot of the 10-fold cross-validation with 5 repeats of the Elastic Net model to predict chronological age using the median-normalized, logarithmically-transformed peptide abundances measured in the full dataset of 15,969 peptides (A). The confusion matrix of this model (B) illustrates that the model performs best in the middle of the age-range tested and is less accurate at predicting chronological age at the earliest and latest ages of the cohort. The q-values of peptides from the peptide-level model are plotted on the y-axis against the q-values proteins from the protein-level model (C). There were 96 peptides that reached significance ($q < 0.01$) in the peptide-level model with proteins that were not significant in the protein-level model.

### Supplementary Materials and Methods

#### *Animal Care and Housing*

C57BL/6J mice of both sexes were sourced from and housed at The Jackson Laboratory (Bar Harbor, ME, USA; Stock Number 000664). The mice were housed in groups of 5 or less and split by sex and maintained on pine shavings in a pathogen-free room on a 12-hour light/dark cycle (6 AM – 6 PM). The mice had *ad libitum* access to both food (LabDiet 5KOG) and acidified water. All experiments complied with the National Institutes of Health Guide for the Care and Use of Laboratory Animals (National Research Council), were approved by The Jackson Laboratory's Animal Care and Use Committee, and were performed in accordance with Animal Research guidelines.

#### *Plasma collection*

Blood was drawn using a submental blood collection protocol and transferred into EDTA blood collection tubes. The plasma collection tubes were spun down at 14,000 rpm (4°C) for 10 minutes. Taking care not to disturb the buffy coat, plasma was drawn off the top, aliquoted in 60 µL aliquots, frozen, and stored at -80°C until processing.

#### *Quality control measures evaluating sample preparation, liquid chromatography-mass spectrometry system suitability, and quantitative results*

Prior to beginning sample preparation, samples and external quality control samples were assigned to one of two batches. The external quality control samples are described below. We used a balanced block design approach similar to published work (Burger et al., 2021; Čuklina et al., 2021) to generate two sample batches of 48 and 56 wells, respectively. We considered the age, sex, and group size of the mice to ensure even distribution of the sample groups across batches. The processing was done in 96-well plates, and the samples were randomized such that there was minimal positional bias in a group on the plates. One of each external quality control sample was included in each row of the plate (10 samples, 2 quality control samples per row). Additionally, a qualitative scoring approach was used for later sample annotation of the degree of hemolysis. A single operator assigned each sample a score of hemolysis falling into the following qualitative bins: Very low, low, moderately low, moderate, moderately severe, severe, and very severe.

We employed a quality control (QC) framework which has been described in detail previously (Tsantilas et al., 2024). Briefly, this included internal QCs to evaluate sample preparation, system suitability methods to track LC-MS function, and external QC samples to assess batch variation and the impact of signal processing on variation. Key reagent sources are listed in Supplementary Table 1. Each experimental and external quality control sample was spiked with 800 ng of an exogenous protein - yeast enolase 1 - to evaluate sample preparation. This included the denaturation, reduction, alkylation, sample clean-up, and tryptic digestion.

Targeted (parallel reaction monitoring) runs of a standardized system suitability sample (600 fmol of digested bovine serum albumin (BSA) and 150 fmol PRTC per injection) were included before, during, and after sample runs. These system suitability runs and the PRTC peptides in samples were evaluated to assess the quality of the chromatography and instrumentation before and during analysis. We analyzed four of these system suitability runs prior to any sample analysis. After every six sample runs, another system suitability run was analyzed.

Two additional pooled plasma samples were generated to be processed and analyzed repeatedly alongside the experimental samples as a “known unknown”. These external QC samples include the inter-batch (IBQC) and inter-experiment (IEQC) external QC samples. The IBQC and IEQC included the same protein (yeast enolase 1) and peptide (PRTC) internal QCs as the experimental samples. The IBQC was a pool of equivalent volumes of plasma collected from 12 fasted, C57BL/6J mice that were part of the experimental mouse cohorts. The mice were evenly split by sex, and their chronological ages spanned 25 to 97 weeks (5.85 - 22.42 months), which was representative of the experimental group. All volumes of individual plasma were pooled, mixed well, aliquoted into 20 µL increments, and stored at -80°C until processing. The IEQC sample was a commercially purchased pool of C57BL/6N mouse plasma that had been previously freeze-thawed and aliquoted prior to generating the 20 µL aliquots that were frozen and stored at -80°C until processing. The protein concentration of the 12 individual mouse samples was determined using a Pierce™ BCA Protein Assay Kits to verify they were suitable for use as a control. The samples were an average of 55.5 ± 4 μg protein/μL relative to a standard curve spanning a range of 2 - 0.025 μg/μL BSA.

***Extracellular vesicle enrichment (Mag-Net), protein digestion, and clean-up***

The Mag-Net protocol (Wu et al., 2025) was used to enrich circulating extracellular vesicles in mouse plasma and digest captured proteins for analysis of the resulting peptides by liquid-chromatography and data-independent mass spectrometry. A Thermo Scientific™ KingFisher Apex was used to run the Mag-Net protocol including particle enrichment, washes, PAC, and digestion. Magnetic beads were moved from plate to plate into different buffer steps.

Briefly, Mag-Net combines a charge and size-based membrane particle capture followed by Protein Aggregation Capture (PAC) and tryptic digestion. Beginning with 20 μL of plasma for each individual mouse, samples were diluted in Dulbecco's phosphate-buffered saline and spiked to 1X of HALT protease and phosphatase inhibitor cocktail. Samples were mixed 1:1 with binding buffer (100 mM Bis-Tris Propane (BTP) and 150 mM sodium chloride, pH = 6.3) and incubated for 10 minutes. The enrichment was performed using 5 μL of MagReSyn® strong anion exchange (SAX) magnetic microparticles (ReSyn Biosciences). This maintained the previously published, optimized bead:plasma ratio for Mag-Net using 25 μL of 20 μg/μL of SAX beads per 100 μL of neat plasma. Beads were equilibrated in 500 μL of wash buffer (50 mM Bis-Tris Propane (BTP) and 150 mM sodium chloride, pH = 6.5) and washed again in 500 μL of wash buffer prior to binding the membrane-bound particles to the MagReSyn® SAX beads. After binding, the sample-bound beads were washed 3 times in 500 μL of wash buffer. The beads were moved into the final well containing 50 mM Tris (pH 8.5), 1% SDS, 10 mM TCEP, and 800 ng of yeast enolase (internal QC). Samples were denatured and reduced for 1 hour at 37°C on the KingFisher. Offline, samples were spiked with 15 mM iodoacetamide and incubated in the dark for 30 minutes to alkylate proteins, and then quenched with 10 mM dithiothreitol. Proteins were precipitated onto the beads by bringing samples to 70% acetonitrile and incubated at room temperature for 10 minutes. Using the KingFisher Apex, five sequential washes in 1 mL of solution were used for sample clean-up: three of 95% acetonitrile and two of 70% ethanol. The washed, precipitated proteins on the MagReSyn® beads were moved to a new plate containing

50 mM Tris (pH = 8.50) containing 1 µg of porcine trypsin and digested for 1 hour at 47°C. Peptides were acidified to 0.48% trifluoroacetic acid.

***Liquid-chromatography and mass spectrometry: Samples***

One µg of each digested sample spiked with 150 femtomole of Pierce Retention Time Calibrant (PRTC) peptides were loaded onto the system.

The Mag-Net enrichment and depletion evaluation from neat plasma were performed using a Thermo Orbitrap Exploris 480. LC separation was done using a 150 µm fused silica Kasil1 (PQ Corporation) fritted microcapillary trap loaded with 3.5ncm of 3 µm Reprosil-Pur C18 (Dr. Maisch) reverse-phase resin coupled with an 75 µm inner diameter picofrit (New Objective) analytical column containing 30 cm of 3 µm Reprosil-Pur C18 attached to a Thermo EASY-nLC 1200. Buffer A was 0.1% formic acid in water and buffer B was 0.1% formic acid in 80% acetonitrile in water. The strong needle wash was 50% acetonitrile in water, and the weak needle was 0.1% trifluoroacetic acid/2% acetonitrile in water. The 60-minute sample LC gradient consisted of 4 to 6% B in 42 seconds, 6 to 7% B in 18 seconds, 7 to 40% in 55 minutes, 40 to 55% B in 1.5 minutes, 50 to 99% B in 18 seconds, concluding with 2.5 minutes at 99% B. Peptides were eluted from the column with a 40°C heated source (CorSolutions) and electrosprayed into a Thermo Orbitrap Exploris 480 Mass Spectrometer with the application of a distal 2.2 kV spray voltage into a 300° C ion transfer tube. Mass spectrometry analysis was split between a chromatogram library and samples. First a chromatogram library of 6 independent injections was analyzed from a pool of the EV samples. For each injection, a cycle of one 30,000 resolution full-scan mass spectrum with a mass range of 110 *m/z* (395-505 *m/z*, 495-605 *m/z*, 595-705 *m/z*, 695-805 *m/z*, 795-905 *m/z*, or 895-1005 *m/z*) followed by a data-independent MS/MS spectra collection at 30,000 resolution, AGC target of 100%, Auto maximum injection time, 27% normalized collision energy with a 4 *m/z* overlapping isolation window. The chromatogram library data was used to quantify proteins from individual sample runs. These individual runs consisted of a cycle of one 30,000 resolution full-scan mass spectrum with a mass range of 395-1005 *m/z*, AGC target of 100%, Auto maximum injection time followed by a data-independent MS/MS spectra collection at 30,000 resolution, AGC target of 100%, Auto maximum injection time, 27% normalized collision energy with an overlapping 12 *m/z* isolation window, and a mass range of 200-2000 *m/z*. Application of the mass spectrometer and LC solvent gradients were controlled by the ThermoFisher Xcalibur data system.

For the aging cohort, data collection was moved to a newer system. We used a 300 µm diameter PepMap™ Neo Trap Cartridge (Thermo) trap loaded with 5 µm diameter C18 particles was coupled with a 150 µm diameter PepSep analytical column packed with 8 cm of 1.5 µm C18 beads (ESI Source Solutions). Separation was performed using a Thermo Vanquish Neo. Buffer A is 0.1% formic acid in water and Buffer B is 0.1% formic acid in 80% acetonitrile. The strong needle wash was 50% acetonitrile and the weak needle wash was 0.1% trifluoroacetic acid in 2% acetonitrile. The 24-minute LC gradient consisted of 3 to 4% B in 0.3 minutes, 4 to 4.5% B in 0.7 minutes, 4.5 to 38% in 21 minutes, 38 to 50% B in 0.5 minutes, 50 to 99% B in 0.5 minutes, concluding with 1 minute at 99% B. Peptides were eluted from the column with a 50°C heated source (CorSolutions) and electrosprayed into a Thermo Orbitrap Astral Mass Spectrometer with the application of a distal 2.5 kV spray voltage into a 300° C ion transfer tube.

***Liquid-chromatography and mass spectrometry: System Suitability***

Targeted parallel reaction monitoring (PRM) runs were included before, during, and after experimental sample for the aging cohort runs to assess the LC-MS system suitability. The target mass list included 17 peptides (all z = 2) which are summarized in Supplementary Table 2. For the Astral, the 10 minute LC gradient was as follows: 0 to 9% B in 1 minute, 9 to 40% B in 7 minutes, 40 to 60% B in 30 seconds, 60 to 99% B in 30 seconds, and 30 seconds at 99% B. A 50 ms precursor scan (400-810 m/z) was followed-up with a set of targeted MS/MS scans (200-1700) where spectra were acquired using the Astral detector, with a standard AGC target of 10,000, a maximum injection time of 35 s, and a 30% normalized collision energy. Peptides were eluted from the column with a 50°C heated source (CorSolutions) and electrosprayed into a Thermo Orbitrap Astral Mass Spectrometer with the application of a distal 2.5 kV spray voltage into a 300° C ion transfer tube. For the Exploris, the 40-minute system suitability gradient consisted of 0 to 16% B in 5 minutes, 16 to 35% in 20 minutes, 35 to 75% B in 5 minutes, 75 to 100% B in 5 minutes, followed by a wash of 9 minutes and a 30-minute column equilibration. The 110-minute sample LC gradient consists of a 2 to 7% for 1 minute, 7 to 14% B in 35 minutes,

14 to 40% B in 55 minutes, 40 to 60% B in 5 minutes, 60 to 98% B in 5 minutes, followed by a 9-minute wash and column equilibration.

On the Exploris for the enrichment/depletion experiment, a cycle of one 120,000 resolution full-scan mass spectrum (350-2000 *m/z*) followed by a data-independent MS/MS spectra on the loop count of 76 data-independent MS/MS spectra using an inclusion list at 15,000 resolution, AGC target of 4e5, 20 millisecond (ms) maximum injection time, 33% normalized collision energy with an 8 *m/z* isolation window. The same settings were used on the Exploris as in the Astral for peptide elution, source temperature, distal spray voltage, and ion transfer tube temperature.

#### *Mass spectrometry signal processing*

Using a Nextflow pipeline (https://github.com/mriffle/nf-skyline-dia-ms), the DIA MS data were converted to mzML format using msConvert (Chambers et al., 2012), peptides identified using EncyclopeDIA, version 2.12.30 (Searle et al., 2018, 2020), q-values and posterior error probabilities at the peptide-spectrum match (PSM) level were acquired using Percolator, version 3.06, (github.com/percolator/percolator/releases) (Käll et al., 2007), and uploaded to Skyline for protein grouping, visualization and data dissemination. Searches were performed using a *Mus musculus* reference proteome FASTA file (Uniprot Proteome ID: UP000000589, downloaded February 2, 2023) appended with the internal QC yeast enolase 1. An earlier version of the Nextflow pipeline was run for the EV enrichment/depletion experiment described in the next section “Enrichment and depletion relative to total plasma” using Nextflow version 23.10.2 and an updated version of Nextflow version 24.04.4 was used for the aging cohort. The raw data, parameter files used in Nextflow DIA analyses, and the processed data downloaded from Skyline used as input to generate Figures 2-5, Figures S1-S5, Tables 1-2, and all Supplementary Tables are available on PanoramaWeb

System suitability and internal controls in the aging cohort samples and external control samples were imported into Skyline version 24.04.4 (Pino et al., 2020).

#### *Enrichment and depletion relative to total plasma*

Three replicate wells containing 100 µL of the C57BL/6N commercial pool used as an IEQC were processed with Mag-Net, denatured, reduced, alkylated, and digested as described in the previous section. Alongside these samples, 1 µL of the same plasma was denatured, reduced, alkylated, and digested as described. The same internal QCs (yeast enolase 1, PRTC peptides) described in the previous section "Liquid-chromatography and mass spectrometry" were included. The data from this experiment was collected using the Nano Easy-LC and a Thermo Orbitrap Exploris 480 method described above.

#### *Particle counting*

The C57BL/6N commercial pool used as an IEQC was also used to demonstrate that Mag-Net could isolate EVs from mouse plasma. Enrichment of membrane-bound particles was done in technical duplicates with 100 µL of starting plasma volume as described above. However, rather than exchanging the wash buffer with SDS and Tris, the particles were eluted from the beads using 100 µL elution buffer of 25 mM Bis Tris Propane (pH = 6.5), 1 M NaCl, 0.1% Tween 20. The particles were diluted 1:25 in water and analyzed using a Nanosight NS300 (Malvern Panalytical Ltd) fitted with a standard gasket (NTA4027) and measured at 25.4 °C ± 0.2. Samples were injected manually. Videos and particle counts for each sample were captured in 5 replicate measurements for each capture using the Nanosight Nanoparticle Tracking Analysis (NTA) software (Malvern Panalytical Ltd, version NTA 3.4, Build 3.4.4).

#### *General peptide and protein-level analysis*

Additional analysis and figure generation was performed outside of Nextflow using R (R version 4.4.1, http://www.r-project.org) run in RStudio (RStudio 2024.09.0+375 "Cranberry Hibiscus" Release) in the format of R Markdown files and Python (Van Rossum & Drake, 2009) with Jupyter Notebooks. Log-transformed median-normalized peak areas were used for analyses of mass spectrometry results performed outside of Skyline unless specifically noted for the purposes of data quality assessment in Supplementary Figure 2.

#### *Spearman correlation and Reactome pathway analysis*

Spearman correlation was performed assuming a two-sided test using the age of the mice and the normalized protein abundance to calculate spearman rho (ρ) and a p-value. The resulting p-values were adjusted using the Benjamini-Hochberg correction (Benjamini & Hochberg, 1995) and used for subsequent p-value cutoffs in Figures 2, 3, S3 and S4. Using the list of proteins positively or negatively correlated proteins by Spearman Correlation ($\rho > 0.3$ or $\rho < -0.3$, adjusted p-value < 0.05), a high-level analysis of implicated pathways was generated in R using packages in the Bioconductor framework. The associated mouse Uniprot Accession numbers were converted into a list of ENTREZ gene IDs using the “clusterProfiler” package (Xu et al., 2024) and the Organism database “org.Mm.eg.db” (Carlson, 2019), version 3.21. The Reactome Pathway Enrichment was performed using the “ReactomePA” package (Yu & He, 2016) with the organism database *Mus musculus.*

#### *Ordinary Least Squares (OLS) of Protein Abundance*

OLS analyses were implemented in Python (Van Rossum & Drake, 2009) using statsmodels (Seabold & Perktold, 2010), NumPy (Harris et al., 2020), scikit-learn (Pedregosa et al., 2011), and SciPy (Virtanen et al., 2020). Raw abundances were recovered by exponentiating log2-transformed intensities, then normalized via natural-log transformation and z-score standardization (scikit-learn StandardScaler). For each feature, an OLS model was fit regressing scaled abundance on age and sex using the statsmodels formula API. P-values for the age coefficient were corrected for multiple testing using the Benjamini–Hochberg false discovery rate (FDR) procedure (Benjamini & Hochberg, 1995), with features having FDR-adjusted q-values < 0.01 considered significant.

#### *OLS of Protein CV*

OLS analyses were implemented as with protein abundance. For each protein, we calculated the mean abundance and CV separately for each of four subgroups defined by age (split into two groups of two) and by sex, then fit a linear regression model with CV as the dependent variable and mean abundance, age group, and sex as independent variables. The following formula was used:

$$\widehat{CV} = \beta_0 + \beta_1 \cdot meanAbundance + \beta_2 \cdot isOld + \beta_3 \cdot isMale$$

Two comparisons were done estimating a threshold corresponding to 70-80% of the median C57BL/6J lifespan and when they are known to be declining in health (Ogiso et al., 2025; Yuan et al., 2009). First younger than 21 months vs. 21 months and older, and then 5-15 months vs. 17-21 months.

#### *Elastic Net Regularized Linear Regression*

An Elastic Net regression model (Zou & Hastie, 2005) was used to predict chronological age from feature abundances. Elastic Net combines L1 (lasso) and L2 (ridge) penalties, enabling simultaneous feature selection and coefficient shrinkage; hyperparameters were set to $\alpha$=0.01 and *l1_ratio=0.3*. Optimal hyperparameters were estimated using Bayesian optimization with the Python library “scikit-learn”. Features were log10-transformed, and sex was included as a binary covariate. Model performance was estimated using repeated *k*-fold cross-validation (10 folds, 5 repeats) implemented with scikit-learn's RepeatedKFold. To prevent data leakage, z-score standardization (scikit-learn StandardScaler) was fit on the training partition and applied to the test partition independently within each fold. Model accuracy was assessed by mean absolute error (MAE) and $R^2$ computed via linear regression of predicted versus true values (SciPy linregress). Robust feature importances were derived by averaging Elastic Net coefficients across all folds and recording the frequency with which each coefficient was non-zero. A final model was constructed using Elastic Net with all features using the same hyperparameters as during cross-validation, and these final coefficients were used to report the number of features kept or removed from the model. All analyses were implemented in Python using scikit-learn, NumPy, and SciPy.

#### ***Data accessibility and figure generation***

All raw files, Skyline documents, processed results used as input for figure generation, FASTA files, EncyclopeDIA files, metadata, Nextflow workflow configuration files and output are available on PanoramaWeb (Murine Plasma EVs). The data input <25 MB, R Markdown files, and Jupyter Notebooks used to generate Figures 2-5, Supplementary Figures 1-5, and Supplementary Tables 1-15 are available on GitHub: (manuscript-aging-mouse-ev). Any input files larger than 25 MB are noted in the ReadMe and freely available on PanoramaWeb.

***Declaration of generative AI and AI-assisted technologies***

During the preparation of this work, Claude (Anthropic) was used for three purposes: to assist the author's in writing portions of the analysis, to write and revise code for figure generation, and to improve the readability and conciseness of the text. In all cases the authors reviewed, edited, and tested the output, and take full responsibility for the content of the publication and for the correctness of the released code.
